# Signals from the bone marrow B cell niches shape pre-leukemic fate in murine B cell acute lymphoblastic leukemia

**DOI:** 10.1101/2025.02.14.638228

**Authors:** Marjorie C Delahaye, Simon Léonard, Jeoffrey Pelletier, Jérôme Destin, Florence Bardin, Mourad Ounis, Céline Monvoisin, Naïs Prade, Laurine Gil, Audrey Dauba, Pierre Milpied, Ahmed Amine Khamlichi, Eric Delabesse, Tony Marchand, Cyril Broccardo, Michel Aurrand-Lions, Bastien Gerby, Stéphane JC Mancini

## Abstract

The bone marrow (BM) microenvironment plays a key role in supporting B cell development. In acute lymphoblastic leukemia (B-ALL), the acquisition of oncogenic driver mutations blocks B cell differentiation at specific stages. When these pre-leukemic cells acquire secondary mutations, B-ALL develops. However, the role of the BM microenvironment in pre-leukemic cell fate remains unknown. Here, using a murine model of spontaneous B-ALL development, we show that disrupted pre-BCR signaling in pre-leukemic cells modifies their fate. Blocking expression of the pre-BCR ligand Galectin-1 by the microenvironment impaired pre-leukemic cell proliferation and leukemia-initiating capacity. Consequently, B-ALL development was delayed, and B-ALL had a more mature phenotype, with cells expressing a BCR. Secondary mutations were also altered by changes to Galectin-1 expression, in its absence mutations almost exclusively affected IL-7R signaling rather than both pre-BCR and IL-7R signaling. These results show that signals from BM niches can directly influence pre-leukemic B cell fate.

## Introduction

B cell acute lymphoblastic leukemia (B-ALL) originates in the bone marrow (BM). It is characterized by arrested B cell differentiation and an accumulation of genetic abnormalities including translocations, aneuploidy, gene deletions and point mutations ^1^. Although less common in adults, B-ALL is the most frequent cancer in children. Its development can be initiated as early as in utero, involving a multi-step process ^2^. The evidence for this process is based on studies of monozygotic twins, with twins sharing the same genomic *TEL::AML1* (aka *ETV6::RUNX1*) fusion developing leukemia several years apart ^3^. These observations not only demonstrate that latent pre-leukemic clones could be present in the BM, but also indicate that the initial genetic anomaly is not sufficient to induce leukemia, secondary oncogenic events are required. This need for a second trigger is further supported by the frequency of *ETV6::RUNX1* translocation detected in newborns, which is over 100-fold higher than the incidence of childhood leukemia ^4^. Among second triggers, the main risk factors for leukemogenesis beyond genetic disorders are environmental stresses such as exposure to ionizing radiation, pesticides, or chemicals causing DNA damage ^5,6^.

Endogenous events can also influence leukemia development. Since the immune system is shaped during the first years of life, coinciding with the peak of incidence of pediatric B-ALL, early studies of B-ALL transformation from the initial oncogenic lesion highlighted the role played by infections ^7,8^. The role of infection has since been modeled in mice. Two mouse models of B-ALL have been developed, one harbors a Pax5 haploinsufficiency, whereas the other expresses the human ETV6::RUNX1 fusion protein. Neither of these models developed B-ALL within two years when maintained in pathogen-free conditions ^9,10^. In contrast, within one month of transfer to a conventional facility where they were exposed to common pathogens, a significant number of individuals developed leukemia. Beyond this short time window and the pathogenic trigger, it remains to be determined how pre-leukemic clones progress toward leukemia.

Studies of various leukemogenesis mouse models reveal that pre-leukemic cells accumulate at early stages of B cell development, before the emergence of secondary genetic alterations and the onset of leukemia ^9–12^. B cell commitment occurs at the pro-B cell stage following expression of the B cell-specific transcription factor PAX5 ^13^. Early B cell subsets then develop in supportive niches in the BM, in contact with growth factor-producing mesenchymal cells ^14^. Pro-B cell proliferation and survival are strongly dependent on the expression of the IL-7 receptor (IL-7R) ^15^, and establish tight contacts with stromal cells expressing IL7 ^16,17^. At this stage, the recombination of genes encoding the immunoglobulin heavy (IgH) chain is initiated. Once a functional IgH chain is produced, the cells differentiate toward the pre-B cell stage with surface expression of a pre-B cell receptor (pre-BCR), composed of the IgH chain, the surrogate light chain (SLC), and the signaling molecules CD79a and CD79b ^18^. Pre-BCR signaling then leads to proliferation and differentiation of pre-B cells ^19^. Although ligand-independent signals have been described ^20^, SLC-mediated pre-BCR binding of Galectin-1 (GAL1) – an s-type lectin expressed by BM stromal cells – is clearly involved in the control of pre-B cell proliferation ^21,22^. Pre-leukemic cell accumulation occurs at these B cell stages, consequently, B cell-specific stromal cell niches may fuel leukemogenesis. In support of this hypothesis, mutations inducing mimicry of IL-7R or pre-BCR signaling are recurrent in B-ALL ^23–25^, suggesting that leukemic cells have a decreased dependence on these niche signals. However, a direct effect of the BM microenvironment on the fate of pre-leukemic cells and the acquisition of secondary mutations has not been formally demonstrated.

In the present study, we took advantage of a murine model expressing the human *PAX5::ELN* fusion gene, identified in B-ALL patients ^26^, that spontaneously develops pre-BCR^+^ pre-B cell leukemia ^11^. Pre-BCR signaling was impaired during leukemogenesis by backcrossing these mice onto a GAL1-deficient background. Analysis of pre-leukemic states by flow cytometry and single cell sequencing revealed that the impaired pre-BCR signaling modified the phenotype and the molecular profile of pre-leukemic cells. Most importantly, leukemic development was delayed and the leukemic sub-type as well as the mutational landscape were modified.

## Results

### Pre-BCR signaling defects impair the proliferative capacity of pre-leukemic cells, modifying their phenotype and activation status

Mice expressing the PAX5::ELN fusion protein identified in B-ALL patients (P5E mice) present an initial pre-leukemic population blocked at the pro-B/pre-B transition which expresses the pre-BCR ^11,27^. Expression of this receptor could be a prerequisite for leukemia development. To test this hypothesis, we crossed P5E mice with RAG2^-/-^ (R2^-/-^) mice (P5ER2^-/-^). R2^-/-^ mice are unable to undergo V(D)J recombination and therefore lack pre-BCR expression ^28^.

The overall proportion of CD19^+^ B cells was significantly increased in In P5ER2^-/-^ and P5E mice as compared to R2^-/-^ and WT mice, respectively (Figure 1A,B). This increase could be attributed to an aberrant CD19^+^B220^lo^ population in the BM of 5-week-old P5E mice, which is also present in P5ER2^-/-^ mice. These cells can be considered pre-leukemic as they are not present in control mice and bear no secondary mutations at this age ^11^. Phenotypically, the B220^lo^ pre-leukemic cell population from P5ER2^-/-^ mice showed an accumulation of CD117^+^ pro-B-like cells, similar to pre-leukemic cells from P5E mice (Figure 1D,E). However, in P5ER2^-/-^ mice, the proportion of B220^lo^ cells in the BM was lower than in P5E mice (Figure 1C). Other differences between the cell populations in the two mouse strains also correlated with expression (or not) of an active pre-BCR. Thus, expression of CD2, induced upon pre-BCR activation ^29^, was increased in CD117^+^ pre-leukemic cells from P5E mice but not in P5ER2^-/-^ mice (Figure 1F). These differences correlated with altered leukemia development profiles between the mouse strains. Among P5E mice, 50% developed leukemia within 22 weeks and more than 80% developed it within 35 weeks, in contrast, less than 25% of P5ER2^-/-^ mice developed the disease over the same period (Figure 1G). These results suggest that pre-BCR expression and activation influence leukemogenesis. However, since RAG proteins can contribute to the acquisition of secondary mutations ^30^, we cannot rule out that the strong reduction in leukemia development observed in P5ER2^-/-^ mice could be related to the absence of RAG2, and independent of the absence of pre-BCR expression.

**Figure 1.**
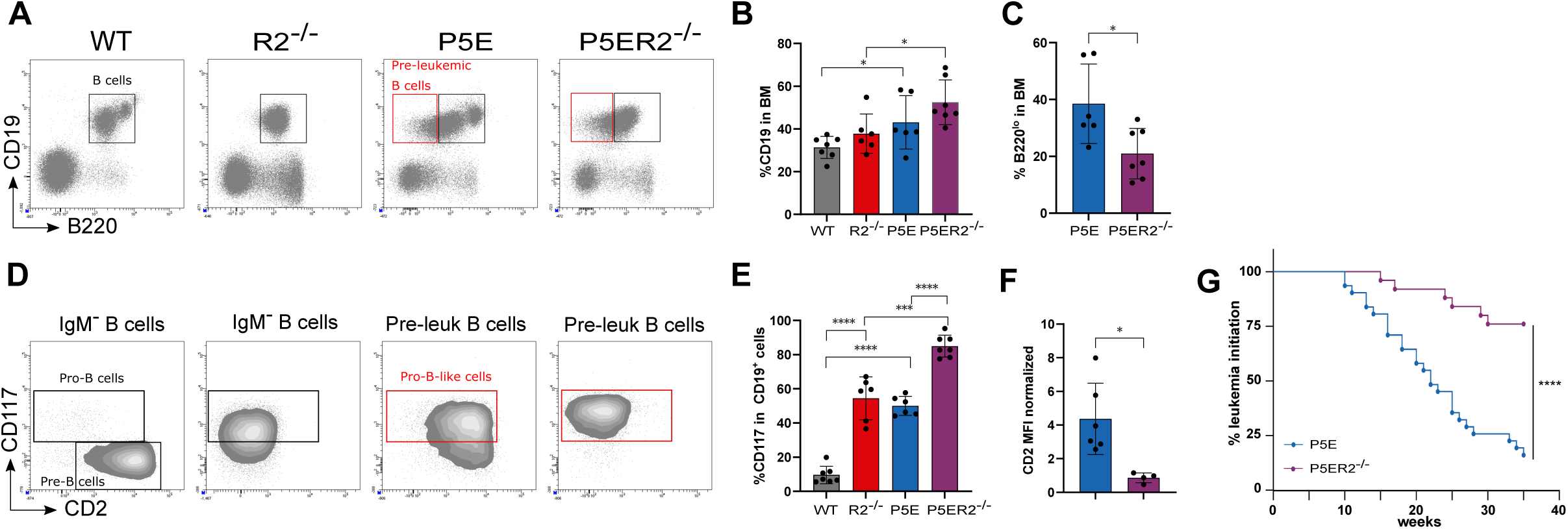
Pre-BCR deficiency impairs leukemogenesis. (**A**) B cells from the BM of 5-7-week-old WT (n=7), R2^-/-^ (n=6), P5E (n=6), and P5ER2^-/-^ (n=7) mice were analyzed by flow cytometry. B cells and pre-leukemic B cells are shown. (**B**) Percentage of total CD19^+^ BM B cells for each genotype. (**C**) Percentage of B220^lo^ pre-leukemic cells. (**D**) Identification of pro-B and pre-B cells among IgM^-^ B cells from WT and R2^-/-^ mice, and of pro-B-like and pre-B-like cells among pre-leukemic B cells from P5E and P5ER2^-/-^ mice, as shown in (A). (**E**) Percentage of CD117^+^ cells among total CD19^+^ cells. (**F**) Normalized mean fluorescence intensity (MFI) for CD2 in pre-leukemic cells from P5E and P5ER2^-/-^ mice. (**G**) Kaplan-Meier survival curve for P5E (n=31) and P5ER2^-/-^ (n=25) mice. Statistical significance was tested using a t-test. *: p-val<0.05; **: p-val<0.01; ***: p-val<0.001; ****: p-val<0.0001.

To distinguish between these effects, we used another mouse strain, deficient for GAL1 expression. GAL1 induces pre-BCR dependent proliferative signaling pathways in both normal and leukemic pre-B cells ^21,31^. To test the influence of GAL1-dependent signaling on the development of pre-BCR^+^ pre-leukemic B cells, we crossed P5E with GAL1-deficient (G1^-/-^) mice (P5EG1^-/-^ mice). A B220^lo^ pre-leukemic population was still present in the absence of GAL1 (Figure 2A, upper panel). However, although the proportion of CD19^+^ B cells was significantly higher in P5EG1^-/-^ as compared to WT or G1^-/-^ mice, it was lower than in P5E mice (Figure 2B). This difference between P5E and P5EG1^-/-^ mice was linked to a decrease in the proportion of pre-leukemic B cells in the BM (Figure 2C). The decrease was not the consequence of escape from the pre-leukemic state, as the proportions of normal B220^int/hi^CD19^+^ pre-B, Immature B, and recirculating B cells were similar in P5E and P5EG1^-/-^ mice (Figure S1A-C). However, the accumulation of CD117^+^ pro-B-like cells among B220^lo^ pre-leukemic cells was lower in the absence of GAL1, with a corresponding increase in numbers of more mature CD2^+^ pre-B-like cells (Figure 2A lower panel, Figure 2D,E). In parallel, PLCγ2 phosphorylation, induced downstream of the pre-BCR, was decreased in P5EG1^-/-^ mice (Figure 2F). Furthermore, analysis of BrdU incorporation 15 h after intra-peritoneal injection demonstrated that, in the absence of GAL1, the pre-leukemic cells had a significantly impaired proliferative capacity (Figure 2G). Finally, and as described previously ^27^, the decreased pre-BCR activation was associated with increased IL-7R signaling, and thus higher STAT5 phosphorylation *in vivo* (Figure 2H).

**Figure 2.**
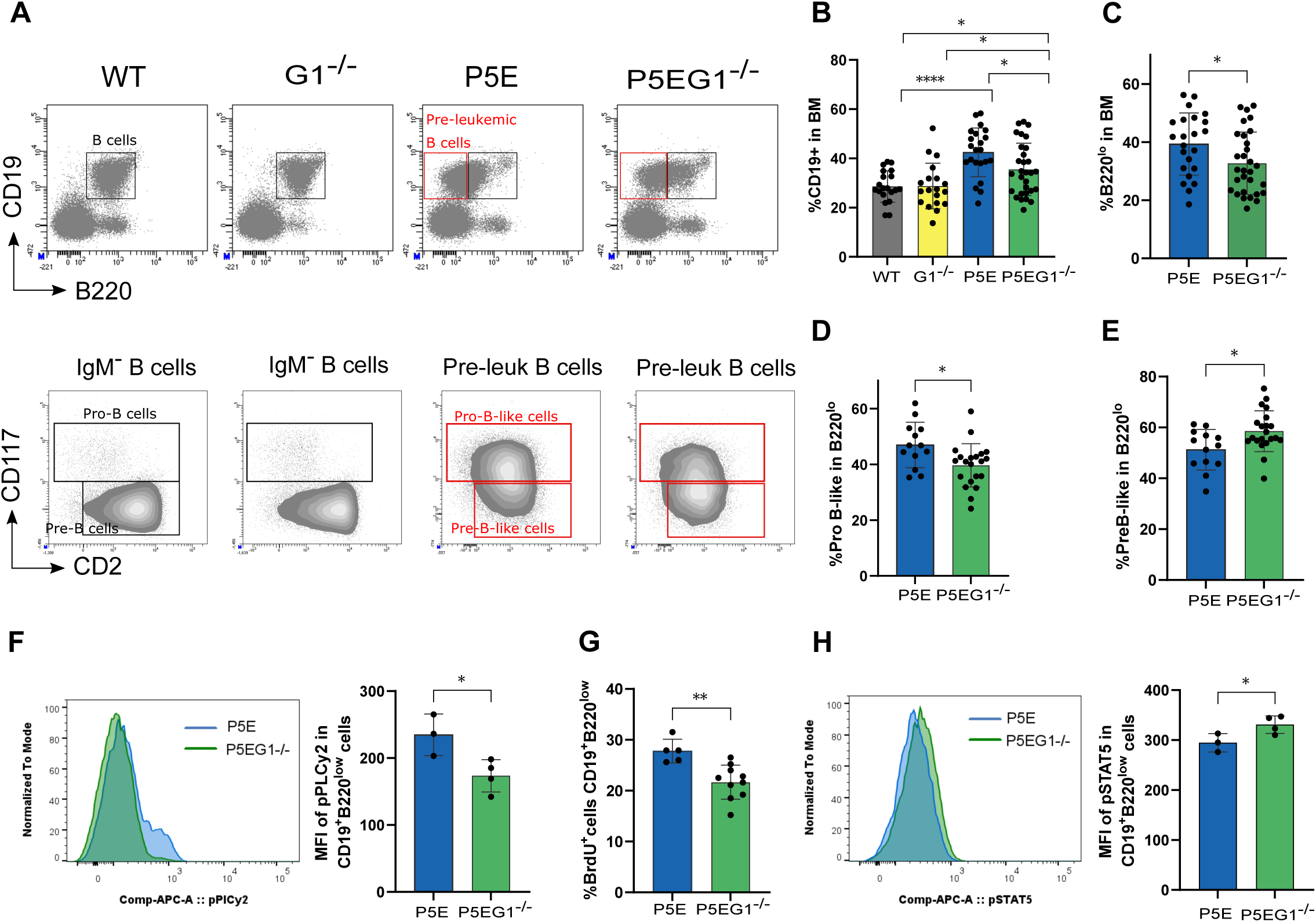
GAL1 stimulates proliferation of pre-leukemic cells and modifies their phenotype and activation status. **A.** B cells from the BM of WT (n=20), G1^-/-^ (n=19), P5E (n=22) and P5EG1^-/-^ (n=30) mice were analyzed by flow cytometry as in Figure 1A,D. Percentage of total CD19^+^ (**B**) and B220^lo^ pre-leukemic cells (**C**). (**D**) Percentage of pro-B-like and (**E**) Pre-B-like cells among B220^lo^ pre-leukemic cells. (**F**) PLCy2 phosphorylation in CD19^+^B220^lo^ pre-leukemic B cells was measured by flow cytometry for 5-7-week-old P5E and P5EG1^-/-^ mice, data are represented as MFI. (**G**) BrdU incorporation by B220^lo^ pre-leukemic cells was determined for 5-7-week-old P5E (n=5) and P5EG1^-/-^ (n=10) mice by flow cytometry performed 15 hours after i.p. injection. (**H**) Phosphorylation of STAT5 was measured by flow cytometry on CD19^+^B220^lo^ pre-leukemic B cells in 5-7-week-old P5E and P5EG1^-/-^ mice, data are expressed as MFI. Representative histograms are shown for (A), (F) and (H). Statistical significance was tested using a t-test. *: p-val<0.05; **: p-val<0.01; ***: p-val<0.001; ****: p-val<0.0001.

Altogether, these results strongly suggest that the absence of GAL1 leads to a loss of pre-BCR signal dependence and to an increase in more differentiated pre-leukemic cells.

### Pre-leukemic cells are heterogeneous and present dysregulated developmental and signaling profiles

The molecular heterogeneity of pre-leukemic cells was subsequently assessed by single cell RNA sequencing (scRNAseq). CD19^+^ B cells were first sorted from P5E and P5EG1^-/-^ mice, WT and G1^-/-^ mice. Their molecular characteristics were then compared. A reduction of dimension using a Uniform Manifold Approximation and Projection (UMAP) representation showed clusters shared by non-transgenic (non-Tg) (WT and G1^-/-^) and Tg animals (P5E and P5EG1^-/-^), but also revealed clusters specific to P5E and P5EG1^-/-^ mice (Figure 3A,B).

**Figure 3.**
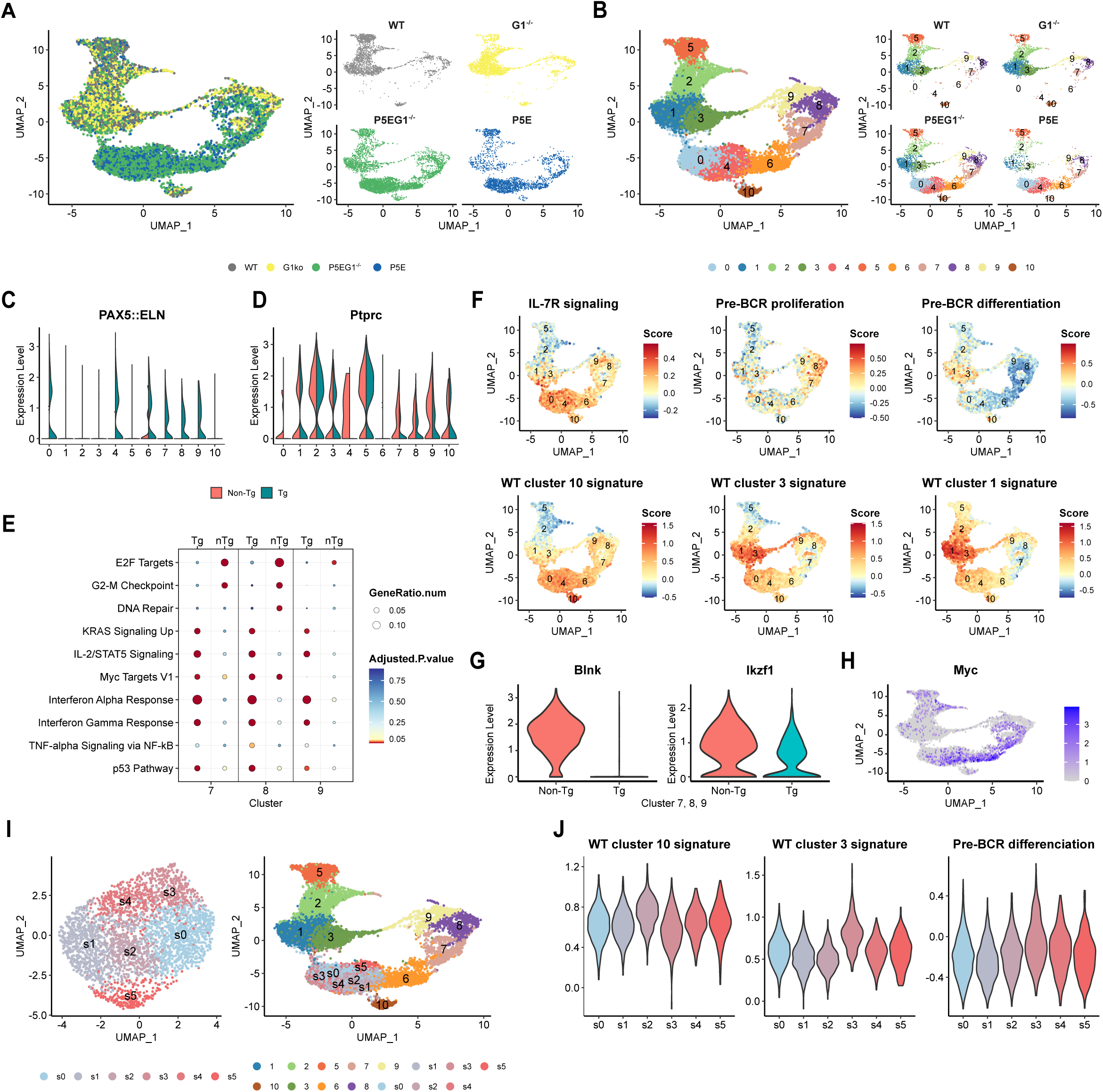
Pre-leukemic cells are heterogeneous and present dysregulated developmental profiles. Total CD19^+^ B cells from 5-week-old WT, G1^-/-^, P5E and P5EG1^-/-^ mice (n=2 for each, except P5EG1^-/-^, n=4) were sorted and analyzed by scRNAseq. Each sample was stained with a specific CD45-conjugated barcode. (**A**) Distribution of B cells in the UMAP space. (**B**) Normal and pre-leukemic B cell heterogeneity were assessed for each genotype, identifying 11 distinct clusters. Expression of the PAX5::ELN chimeric gene (**C**) and of the *Ptprc* gene coding for CD45/CD45R (**D**), are shown for each cluster for non-transgenic (WT and G1^-/-^) and transgenic (P5E and P5EG1^-/-^) samples. (**E**) Differentially expressed genes (DEG) with an adjusted p-value <0.05 between non-Tg and Tg mice were identified for clusters 7, 8, and 9, corresponding to large pre-B cells. The enrichments obtained with MSigDB were assessed for the genes upregulated in both non-Tg and Tg cells. (**F**) WT cells were used to identify the 50 DEG with the highest adjusted p-value for the pro-B stage (cluster 10) and the small pre-B stage (clusters 1 and 3) as compared to all the other clusters. A score was generated for these DEG and for genes representative of IL-7R signaling, pre-BCR proliferation signals, and pre-BCR differentiation signals. The scores were mapped on the UMAP for all cells. (**G**) Violin plots representing the expression of *Blnk* and *Ikzf1* in cells from clusters 7, 8, 9 in non-Tg vs. Tg mice. (**H**) Mapping the expression of *Myc* on the UMAP. (**I**) Subclustering of clusters 0 and 4 extracted from the main UMAP in (B). The six subclusters obtained (left panel) were mapped in the space of the main UMAP (right panel). (**J**) Violin plots representing the scores obtained from the pro-B, small pre-B, and pre-BCR differentiation signatures for each subcluster, as shown in (F).

As shown previously, the differentiation profiles of WT and G1^-/-^ mice at steady state were very similar ^21^. Thus, normal B cell subsets were identified in non-Tg mice based on gene signatures characteristic of the various BM B cell subsets, linked to IL-7R and pre-BCR signaling, and to the cell cycle (Figure S2A,B). The high IL-7R signaling signature confirmed that cluster 10 represented pro-B cells. Clusters 7, 8, and 9 corresponded to large pre-B cells in the G1/S, S/G2M and G2-M phases of the cell cycle, respectively (Figure S2C). As expected, these clusters showed the highest scores for pre-BCR proliferative signals. Furthermore, the switch to pre-BCR differentiating signals, characteristic of the large to small pre-B transition, was already detectable at the extreme end of cluster 9, oriented toward the profile of small pre-B cells in clusters 3 and 1. These two small pre-B cell clusters (Figure S2D) confirmed observations from a previous study ^32^, regarding the existence of an *Iglc*^lo^ and an *Iglc*^+^ cluster (Figure S2D). Finally, clusters 2 and 5 were identified as immature and recirculating B cells, respectively. However, the non-Tg mice were not identical, as the analysis of differentially expressed genes (DEG) between WT and G1^-/-^ cells present in the large pre-B cell clusters showed WT cells were enriched for genes involved in proliferation (Figure S2E), confirming the role of GAL1 in the induction of pre-BCR proliferative signals.

A similar analysis of the Tg mice, P5E and P5EG1^-/-^, revealed the presence of three additional clusters compared to WT and G1^-/-^ mice (clusters 0, 4 and 6; Figure 3B). The polyclonal (pre-)BCR repertoire in all the Tg mice analyzed confirmed the pre-leukemic status of these clusters (Figure S2F). Although the large pre-B cell clusters (7, 8 and 9) were common to both non-Tg and Tg mice, a clear distinction in cell positioning emerged depending on their genotype (Figure 3B). Furthermore, as expected ^27^, the *PAX5::ELN* chimeric gene was not expressed in non-Tg clusters, but only in pre-leukemic clusters 0, 4 and 6, as well as 7, 8 and 9 for Tg samples (Figure 3C). Finally, and in agreement with the flow cytometry-identified decreased expression of B220 (aka CD45R, splice variant of CD45 encoded by *Ptprc*), *Ptprc* expression was decreased in clusters 7, 8 and 9 in Tg mice, confirming that they represent pre-leukemic cells (Figure 3D).

The analysis of DEG between non-Tg and Tg mice in the large pre-B clusters showed that, in contrast to the enrichment for genes associated with proliferation and (pre-)BCR signaling seen in non-Tg animals, in Tg animals, these clusters were enriched for genes related to interleukin signaling and inflammation (Figure 3E). These results were confirmed by analyzing IL-7R and pre-BCR signatures in the large pre-B clusters, revealing an increase in the signature for IL-7R signaling. This increase was counterbalanced by a decrease in the pre-BCR differentiation signature (compare Figure 3F and Figure S2B), as observed previously in P5E mice ^27^. The decreased capacity of large pre-B-like pre-leukemic cells in Tg mice to deliver pre-BCR differentiation signals is most likely due to a loss of expression of the tumor suppressors *Blnk* and *Ikzf1* (Figure 3G), which are involved in the switch from proliferation to differentiation signals ^19,33,34^.

Based on signatures of pro-B and IL-7R signaling, the analysis of clusters 0, 4, and 6 showed a proximity to pro-B cells. These cells were classified as being in the G1 phase of cell cycle (Figure 3B and Figure S2C). In addition, an inter-cluster comparative analysis of DEG revealed a predominance of interleukin signaling and inflammation (Figure S2G). Cluster 6, which is adjacent to cluster 7 (G1/S large pre-B), expressed high levels of *Myc* (Figure 3H) and, as a consequence, was strongly enriched for *Myc* target genes (Figure S2G). This expression pattern is consistent with previous studies reporting that the transition from pro-B to large pre-B cells relies on strong, transient expression of MYC ^35^. Finally, cluster 0 was interesting in that it presented signatures of small pre-B and pre-BCR differentiation. These signatures correlated with the close proximity to and connections between cluster 0 and cluster 1 and 3, the small pre-B clusters (Figure 3F).

To better resolve the heterogeneity of clusters 0 and 4, which both had a pro-B signature in addition to the mixed pro-B/small pre-B signature of cluster 0, we performed subclustering. Six distinct subclusters of pre-leukemic cells were defined that could be linked back to the main UMAP. Subclusters s1 and s5 were adjacent to cluster 6, and subcluster s3 was next to the small pre-B cell cluster 3 (Figure 3I). DEG analysis of the subclusters revealed that subcluster s1 was enriched for elongation and MYC signaling, characteristics also observed in the adjacent cluster 6 (Figure S2H). In contrast, subcluster s5 was mainly enriched for inflammatory signatures. Enrichment for IL-7 signaling was mainly associated with subcluster s3, along with enrichment for BCR signaling that correlates with increased pre-BCR differentiation signals and small pre-B signature (Figure S2H and Figure 3J). This increased BCR signaling signature is particularly revealed by a strong upregulation of *Blnk*, observed only in subcluster s3.

These results strongly suggest that pre-leukemic cells can follow two distinct fates, both linked to IL-7 signaling, but distinguished based on pre-BCR expression. One leads to the emergence of large pre-B-like cells, associated with a combination of pre-BCR proliferative and IL-7R signals, and the other produces small pre-B-like cells, driven by IL-7R signals and pre-BCR differentiation signals.

Based on these results, pre-leukemic cells in P5E mice are heterogeneous and present dysregulated profiles of differentiating B cells blocked between the pro-B, large-pre-B, and small pre-B cell stages of development.

### GAL1 in the pre-leukemic niche sustains pre-BCR proliferative signals and leukemia-initiating cell activity

As small pre-B cells were present in equivalent proportions in P5E and P5EG1^-/-^ mice (Figure S1), we used their numbers to normalize pre-leukemic cell numbers in the two mouse strains. Following normalization, scRNAseq data confirmed pre-leukemic cell numbers were decreased in all P5EG1^-/-^ samples as compared to P5E (Figure 4A). Large pre-B-like clusters and those oriented toward them, and thus toward entry into the cell cycle (cl. 1 and 6), were most affected (Figure 4B). To better assess differences in pre-leukemic development in the presence and absence of GAL1, a pseudo-time analysis was performed using Monocle3, taking pro-B cell cluster 10 as root. Trajectory inference revealed two diverging branches that reconverge in the small pre-B clusters (Figure 4C). Comparing P5E and P5EG1^-/-^ mice on the branch pointing toward large pre-B cells (blue branch) revealed differential expression of genes involved in proliferation, with lower expression levels or faster downregulation in P5EG1^-/-^ (Figure 4D). Genes upregulated at the pre-B cell stage such as *Hspa5* or *Hsp90b1*, implicated in IgH folding and the unfolded protein response ^36,37^, *Sdc4* ^38^ and *Spp1*, upregulated downstream of the BCR ^39^, were also impaired in P5EG1^-/-^ mice. Furthermore, *Vpreb1*, which is part of the pre-BCR, was downregulated earlier than in P5E animals. This downregulation was followed by an upregulation of genes involved in the switch from proliferation to differentiation signals (*Blnk, Ikzf1, Rag1, Il2ra, Cd2, Btg2, Jun*) ^33,34,40–44^. Genes involved in differentiation were also overexpressed in the path pointing toward small pre-B cells (red branch, Figure 4C,D), thus confirming that GAL1 triggers pre-BCR-associated proliferative signals at the expense of ligand-independent pre-BCR differentiation signals. Our previous work with the P5E mouse model showed that, in addition to the pool of proliferative pre-leukemic cells, some cells were in a quiescent state with a capacity to sustain leukemia-initiating cell activity ^27^. We used our previous DEG from RNAseq data of proliferative vs. quiescent P5E pre-leukemic cells to generate scores which we then projected onto the scRNAseq dataset (see Material and Methods). As expected, most of the proliferative cells were found in clusters corresponding to large pre-B cells (Figure 4E left panel), and their relative number was decreased in the absence of GAL1 (Figure 4F). Quiescent cells were distributed over the other pre-leukemic clusters, with the highest proportion located in subcluster s2 (Figure 4E right panel and Figure S2I). In P5EG1^-/-^ mice, the proportion of quiescent cells was most decreased in this subcluster compared to in P5E mice (Figure S2I). In addition, relative numbers of quiescent cells were lower for P5EG1^-/-^ mice in subclusters s1 and s2, and in cluster 6, all of which are oriented toward a large pre-B-like cell fate (Figure 4G). Altogether, these results indicate that quiescence of PAX5::ELN^+^ pre-leukemic cells is decreased in the absence of GAL1, hindering progression toward the proliferative large pre-B-like cell stage.

**Figure 4.**
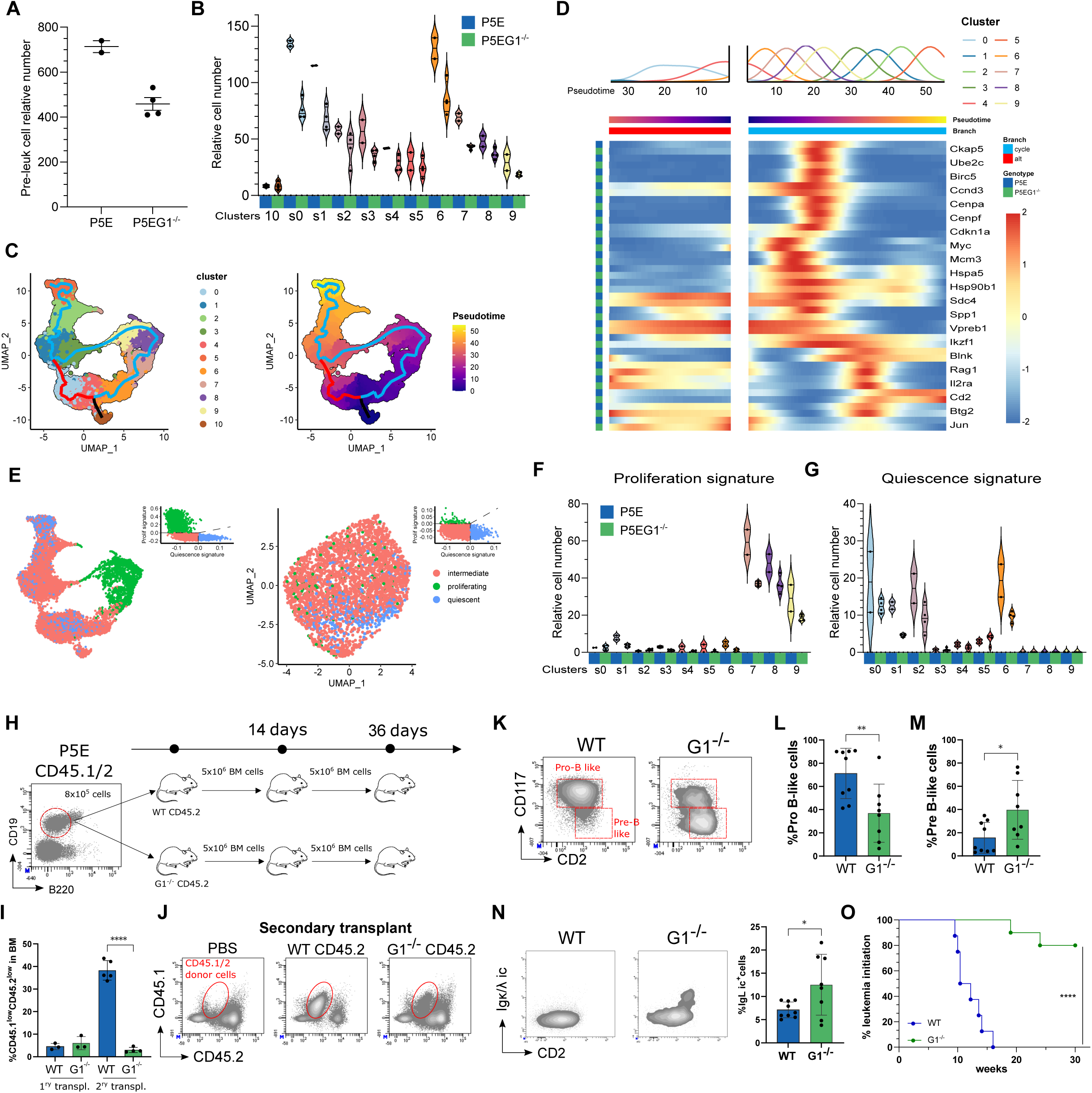
Leukemia-initiating activity of pre-leukemic cells is altered in the absence of GAL1. **(A)** Relative numbers of pre-leukemic cells in P5E and P5EG1^-/-^ mice in the scRNaseq dataset normalized relative to small pre-B cells (Cluster 1+3). (**B**) Relative cell numbers in the pro-B (cluster 10) and the pre-leukemic clusters and subclusters in P5E and P5EG1^-/-^ mice. (**C**) Pseudo-time analysis performed with Monocle3. The trajectory inference is shown with the clusters (left panel) and with the pseudo-time (right panel). The trajectory inference shows two diverging branches starting from the pro-B cell as root, one branch goes toward small pre-B cells (red branch) while the other goes toward large pre-B cells (blue branch). (**D**) Heatmap of the genes varying most between P5E and P5EG1^-/-^ cells along the two branches of the pseudo-time analysis. The frequency of cells in each cluster of the main UMAP are shown above the heatmap. (**E**) Quiescence and proliferation status of each cell in the UMAP space, based on the score generated with the quiescent and proliferative signatures of P5E pre-leukemic cells obtained from bulk RNAseq data. The quiescence or proliferation score for each cell is shown in the upper right of the panels. Violin plots representing the relative number of cells classified as quiescent (**F**) or proliferative (**G**) in each cluster for P5E vs. P5EG1^-/-^ cells. (**H**) Experimental procedure followed for the serial transplantation of pre-leukemic cells sorted from P5E mice and engrafted either in WT or in G1^-/-^ mice. (**I**) Percentage of CD45.1^lo^CD45.2^lo^ pre-leukemic cell engraftment in the BM 14 days after the first transplantation in WT and G1^-/-^ mice, and 22 days after the second transplantation of WT and G1^-/-^ mice. (**J**) Representative flow cytometry analysis of secondary transplants showing CD45.1 and CD45.2 expression. Donor cells are circled in red. (**K)** Representative flow cytometry analysis of CD117 and CD2 expression by pre-leukemic cells. The gating for pro-B-like and pre-B like cells is shown. (**L**) Percentage of pro-B-like cells and (**M**) pre-B-like cells. (**N**) Representative flow cytometry analysis of intracellular expression of IgL (Igκ^+^Igλ) and surface expression of CD2 in pre-leukemic cells, and percentage of intracellular IgL^+^ cells. (**O**) Kaplan-Meier curve representing leukemia development following the third transplantation in WT (n=8) or G1^-/-^ (n=10) mice. Statistical significance was tested using a t-test. *: p-val<0.05; **: p-val<0.01; ***: p-val<0.001; ****: p-val<0.0001.

To test this hypothesis, CD19^+^B220^lo^ pre-leukemic cells from 5-week-old P5E mice with a CD45.1/CD45.2 haplotype were sorted and serially transplanted into CD45.2^+^ WT or G1^-/-^ mice (Figure 4H). Analysis of the first transplant after 15 days showed a similar proportion of pre-leukemic cells in both host environments (Figure 4I). However, while the proportion of pre-leukemic cells increased sharply 3 weeks after the secondary transplant to WT mice (as observed previously ^27^), pre-leukemic cells did not expand in G1^-/-^ hosts following secondary transplants (Figure 4I,J). In the WT background, pre-leukemic cells were mainly blocked at the CD117^+^CD2^lo^ pro-B/large pre-B transition, whereas in a GAL1-deficient background, the majority of pre-leukemic cells were more mature, with a CD117^-^CD2^+^ small pre-B cell phenotype (Figure 4K-M). Furthermore, in the GAL1-deficient background, a larger proportion of pre-leukemic cells expressed an intracellular IgL chain, confirming that their phenotype is more mature than pre-leukemic cells grown in a WT microenvironment (Figure 4N).

Finally, to determine whether pre-leukemic cells grown in a GAL1-deficient microenvironment retained leukemia-initiating cell properties, secondary transplants in WT and G1^-/-^ mice were engrafted to WT and G1^-/-^ tertiary recipients, respectively. Cell fate was monitored until the onset of the first symptoms of leukemia. Among the WT animals, all individuals developed B-ALL between 10 and 16 weeks of transplantation, whereas only 2 out of 10 GAL1-deficient mice developed leukemia (Figure 4O). Taken together, these results demonstrate that in the absence of efficient GAL1-dependent pre-BCR signaling, pre-leukemic cells expressing the PAX5::ELN fusion protein have a more mature phenotype and diminished leukemia-initiating properties. The serial transplantation assay confirmed that the dependence of pre-leukemic cells on GAL1 was not cell-intrinsic, but that it relied on the production of GAL1 by the BM microenvironment.

### Leukemogenesis is delayed and leukemic fate is modified in the absence of GAL1

Since the absence of GAL1 alters the activation status of pre-leukemic cells as well as their molecular and phenotypic profiles, we sought to determine whether these modifications affected B-ALL development. Leukemia development was significantly delayed in P5EG1^-/-^ mice, albeit to a lower extent than in P5ER2^-/-^ mice. Median survival for P5EG1^-/-^ mice was 28 weeks, compared to 22 weeks in P5E animals (Figure 5A). Furthermore, monitoring for circulating leukemic cells confirmed that their emergence was delayed in P5EG1^-/-^ mice compared to P5E mice, but that their growth kinetics were similar (Figure 5B). This observation suggests that the delayed emergence was most likely due to a delay in the acquisition of secondary mutational events. Furthermore, while 85% of B-ALL in P5E mice express a pre-BCR (IgM^+^IgL^-^), in the absence of GAL1 a significant decrease in pre-BCR^+^ frequency is observed, compensated for by the proportion of BCR^+^ cells (IgM^+^IgL^+^) (Figure 5C-E).

**Figure 5.**
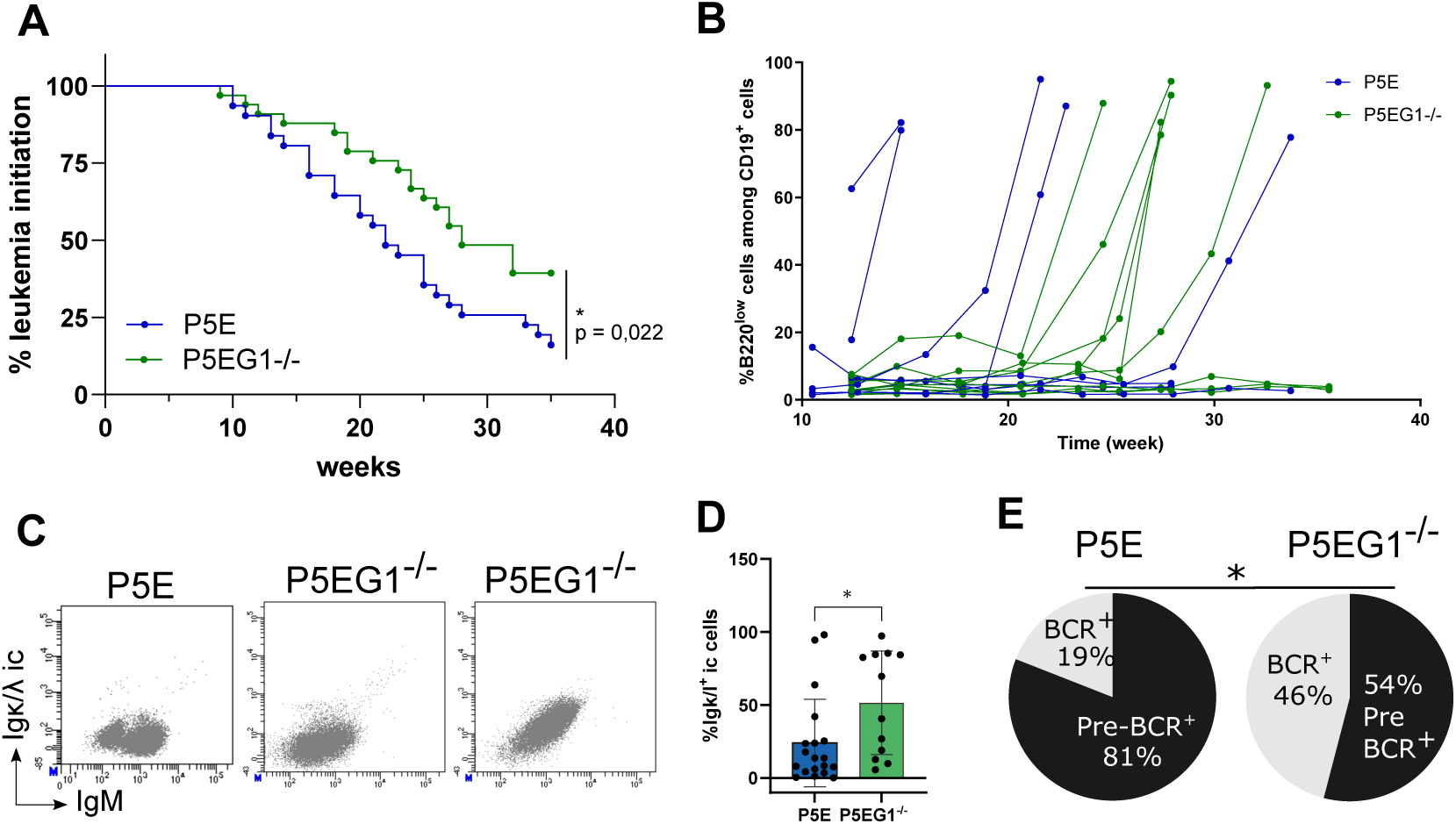
Leukemogenesis is delayed and leukemic fate is modified in the absence of GAL1. (**A**) Kaplan-Meier curve representing leukemia development in P5E (n=31) and P5EG1^-/-^ (n=37) mice. (**B**) Monitoring of the percentage of B220^lo^CD19^+^ (pre-)leukemic cells in the blood of P5E (n=9) and P5EG1^-/-^ (n=10) mice. (**C**) Representative flow cytometry analysis of IgM and intracellular Igκ/λ expression in B-ALL cells from P5E and P5EG1^-/-^ mice. (**D**) Percentage of Igκ/λ expression in B-ALL developing from P5E (n=19) and P5EG1^-/-^ (n=12) mice. (**E**) Percentage of pre-BCR^+^ and BCR^+^ B-ALL developing from P5E (n=35) and P5EG1^-/-^ (n=37) mice. Statistical significance was tested using a t-test (D) or a Fisher’s exact test (E). *: p-val<0.05; **: p-val<0.01; ***: p-val<0.001; ****: p-val<0.0001.

Altogether, these results demonstrate that, through GAL1-dependent pre-BCR signals, the BM microenvironment can influence the leukemic B cell fate.

### Leukemic cells from P5E and P5EG1^-/-^ mice share strong similarities with pre-leukemic cells

To assess the transcriptomic evolution of leukemic cells compared to their pre-leukemic counterparts, we performed scRNAseq on three pre-BCR^+^ B-ALL samples from P5E mice, as well as one pre-BCR^+^ and two BCR^+^ B-ALL samples from P5EG1^-/-^ mice. In addition, CD19^+^ cells from a 5-week-old P5E pre-leukemic mouse were sorted as a control to enhance the integration with the pre-leukemic samples from our first dataset. Repertoire analysis confirmed that, unlike the pre-leukemic sample, the six leukemic samples presented a dominant clone with a unique IgH (Figure S3A). Furthermore, and as expected, the cells with dominant IgH clonotypes among the BCR^+^ P5EG1^-/-^ B-ALL samples were also mostly monoclonal at the level of their rearranged IgL genes. In contrast, IgL clonotypes were diverse and only rarely present in pre-BCR^+^ leukemic cells (Figure S3B,C). Although slightly decreased for one sample, genetic expression of the λ5 and VpreB chains of the SLC, which associates with the IgH chain to form the pre-BCR, remained high in BCR^+^ P5EG1^-/-^ B-ALL samples (Figure S3D). As the SLC is not expressed in B cells at the immature BCR^+^ cell stage, this result confirmed that BCR^+^ leukemic cells are blocked at the late small pre-B cell stage – at the pre-B/Immature B cell transition – as is the case for pre-leukemic cells in P5EG1^-/-^ mice.

After integration of data for leukemic cells (cells expressing the dominant IgH clonotype) to the data for pre-leukemic cells in the first dataset, the cells from the different samples were distributed into 14 clusters, irrespective of their leukemic or pre-leukemic status (Figure 6A and Figure S3E). Importantly, marker genes from the pre-leukemic clustering and subclustering highlighted a similar spatial localization between the integrated leukemic UMAP and the UMAP initially generated for cells from control and pre-leukemic animals (Figure S3F).

**Figure 6.**
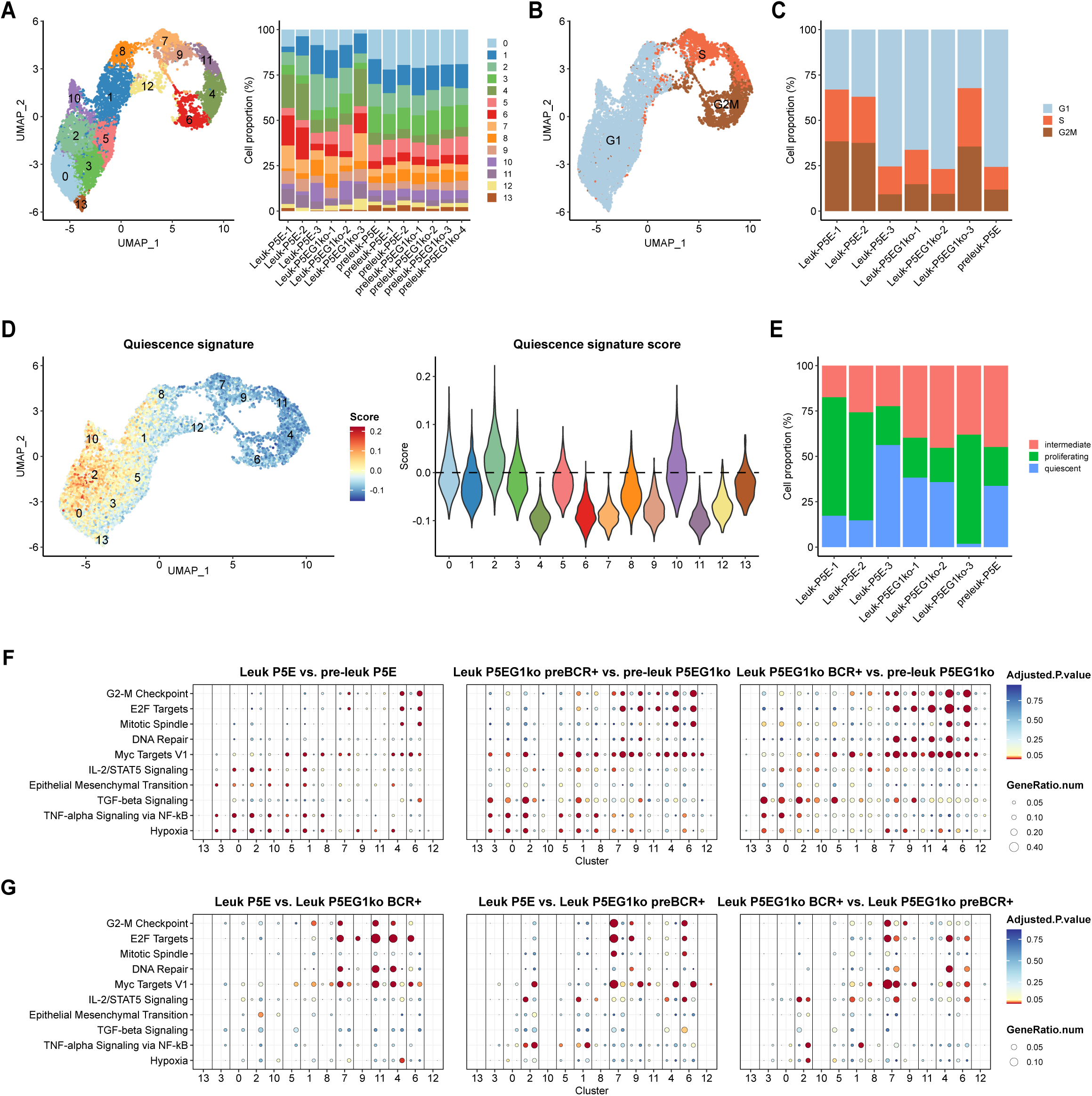
Leukemic cells from P5E and PEG1^-/-^ mice share strong similarities with pre-leukemic cells. CD19^+^ pre-BCR^+^ B-ALL from P5E and P5EG1^-/-^ mice as well as BCR^+^ B-ALL from P5EG1^-/-^ mice were sorted from the BM and analyzed by scRNAseq. CD19^+^ B cells from a pre-leukemic 5-week-old P5E mouse were used as a control. The cells from this dataset and pre-leukemic cells from the dataset used in Figure 3 were integrated, and expression levels normalized before performing another UMAP reduction of dimension. B-ALL cells were selected on the basis of expression of the dominant IgH clonotype. (**A**) UMAP showing the distinct (pre-)leukemic clusters. (**B**) Each cell on the UMAP was tagged with its cell cycle signature by defining a score based on genes characteristic of the S and G2M phases. (**C**) Proportion of cells in the G1, S, and G2M phases of the cycle in each sample. (**D**) Quiescence score for each cell as defined in Figure 4, shown in the UMAP space (left panel) or as violin plots (right panel). (**E**) Proportion of cells tagged as “quiescent”, “intermediate” and “proliferative” in each pre-leukemic cluster, based on their genotype (P5E or P5EG1^-/-^). (**F**) and (**G**) Dot plot of MSigDB enrichments obtained from intra-cluster DEG with adjusted p-values <0.05 in the conditions indicated in the panels.

Cell cycle analysis showed that clusters 7, 9, 11, 4, 6, and part of cluster 12 corresponded to cells in the S and G2M phases of the cycle (Figure 6B). The leukemic samples P5E-1, P5E-2, and P5EG1-3 were mainly cycling, whereas the others showed a distribution very similar to the pre-leukemic samples, whether GAL1 was present in their microenvironment or not (Figure 6C). Cells exhibiting a signature of quiescence were predominantly present in cluster 2, which is related to subcluster s2 from the first dataset (Figure 6D, Figure S2I and Figure S3F). These cells were present in all samples, albeit in lower proportions in samples that had a higher proliferation rate, again whether GAL1 was present or absent (Figure 6E).

At the transcriptomic level, all the leukemic samples overexpressed *S100a4* and *S100a6*, genes implicated in cancer progression (Figure S3G) ^45,46^. DEG between pre-BCR^+^ B-ALL from P5E and P5E pre-leukemic cells or between pre-BCR^+^ or BCR^+^ B-ALL from P5EG1^-/-^ mice and P5EG1^-/-^ pre-leukemic cells revealed a decreased enrichment in ‘E2F targets’, ‘G2-M checkpoint’ or ‘DNA repair’ genes in leukemic compared to pre-leukemic cells (Figure 6F). This decrease may be related to the increase in hypoxia denoted by the enrichment of genes associated with this term, as well as to an attenuated dependence on GAL1-induced pre-BCR signals, since pre-BCR^+^ B-ALL from P5E mice were enriched for genes linked to ‘E2F targets’, ‘G2-M checkpoint’ or ‘DNA repair’ in comparison to both pre-BCR^+^ and BCR^+^ B-ALL from P5EG1^-/-^ mice (Figure 6F,G). MYC and STAT5 signaling, which was initially observed to be enriched in pre-leukemic large pre-B-like cells compared to normal large pre-B cells, was also enriched in leukemic compared to pre-leukemic cells, particularly for clusters in the G0/G1 phase (Figure 3E and Figure 6F). Finally, TGFβ and TNF signaling were enriched in leukemic cells. This signature of chronic inflammation is characteristic of cancer cells.

Therefore, although the individual leukemic samples shared typical features with tumor cells, and despite their distinct phenotypic characteristics, B-ALL retained the heterogeneity observed for pre-leukemic cells, sharing strong similarities at the transcriptomic level with the pro-B/pre-B stages of development.

### Cues from the BM microenvironment influence the acquisition of secondary mutations during leukemogenesis

B-ALL develops in P5E mice by accumulating secondary mutations, mainly in *Ptpn11* and less frequently in *Kras* ^11^, which is activated downstream of the pre-BCR and involved in the proliferative phase of pre-BCR activation ^19^, or in *Jak3*, which is activated downstream of the IL-7R. Targeted sequencing confirmed these mutation patterns and further showed that mutations in the pre-BCR and the IL-7R signaling pathways are mostly mutually exclusive (Figure 7A,B), as described elsewhere ^23^. Therefore, samples in which mutations in both pathways are detected likely indicate the presence of multiple B-ALL clones. Whole exome sequencing analysis of samples that were negative by targeted sequencing identified mutations of genes downstream of the IL-7R, such as gain-of-function mutations of *Jak1* or *Kmt2d* ^47^, and the insertion of a STOP codon at the very beginning of the *Npm1* gene, a negative regulator of STAT5 ^48^ (Figure 7A). In one sample, a mutation in *Shank3*, which negatively modulates ERK signaling ^49^, was also found. As *Shank3* mutations mostly correspond to loss-of-function mutations ^50^, it is likely that this mutation leads to overactivation of the ERK pathway. Given the pivotal role of the IL-7R and the pre-BCR in delivering co-signals to large pre-B cells through the cooperative activation of the ERK signaling pathway leading to the enforced activation of MYC ^35,51,52^, it is possible that the strong IL-7R signaling observed in pre-leukemic cells (compared to WT cells) led to selection of mutations of the ERK pathway ^27^.

**Figure 7.**
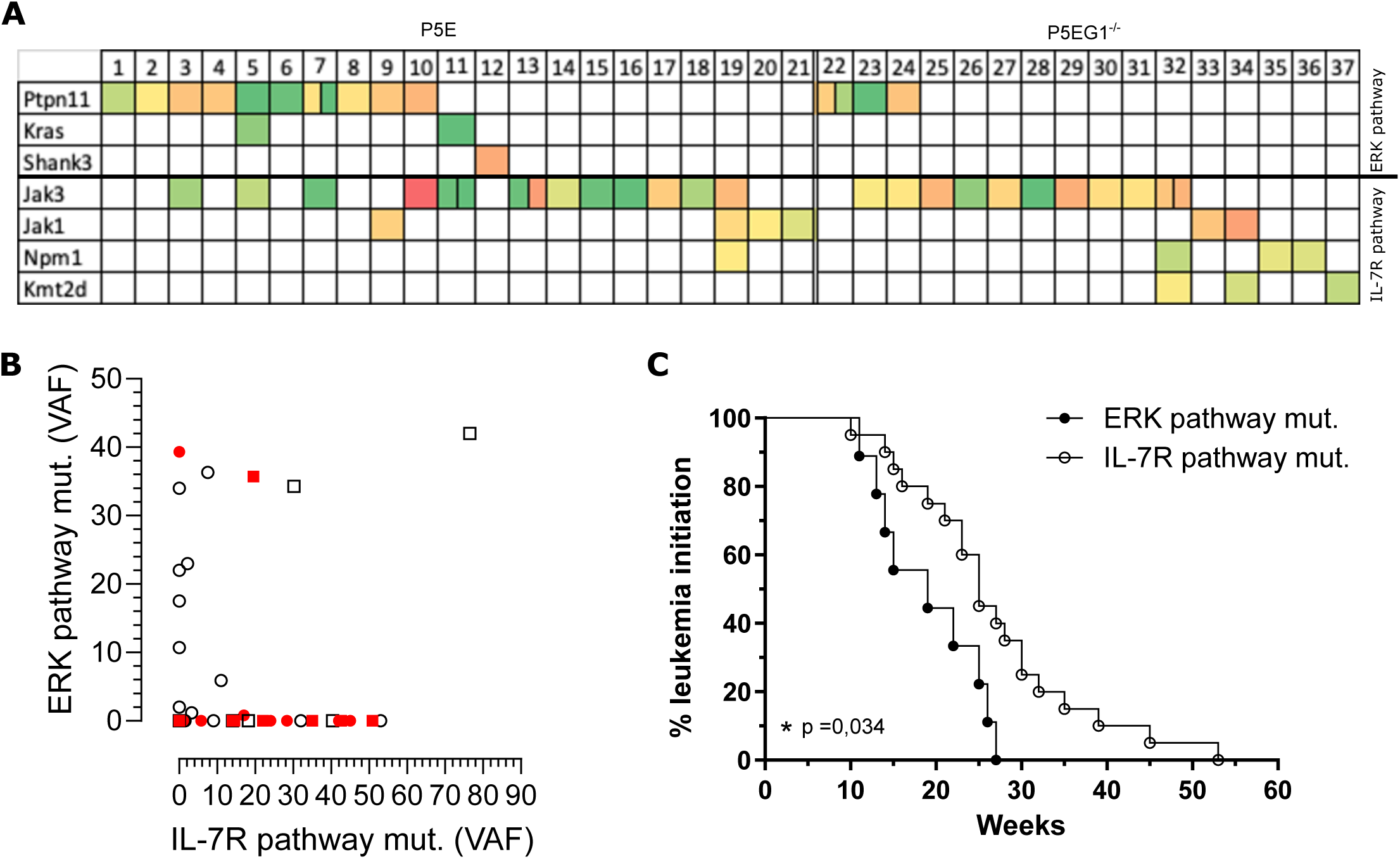
Cues from the BM microenvironment influence the acquisition of secondary mutations during leukemogenesis. (**A**) DNA from the BM of P5E (n=21) and P5EG1^-/-^ (n=16) leukemic mice was analyzed by targeted sequencing and whole exome sequencing (WES). Each row represents a leukemic sample, each line represents mutated genes. The heatmap represents the variant allelic frequency (VAF) from 1% (green) to 77% (red). (**B**) Dot plot showing the VAF for mutated genes downstream of the pre-BCR and downstream of the IL-7R for samples obtained from P5E (white dots) and P5EG1^-/-^ (red dots) mice. Circles and squares represent data obtained from targeted sequencing and WES, respectively. (**C**) Kaplan-Meier curves representing the emergence of mutations mimicking activation of the pre-BCR (n=9) and the IL-7R (n=20).

Analysis of B-ALL from P5EG1^-/-^ mice revealed a limited number of samples bearing mutations in the ERK pathway, represented only by *Ptpn11* mutations, but many mutations downstream of the IL-7R (Figure 7A,B). These results confirm that ERK mutations are induced following activation of the pre-BCR, and that this activation is GAL1-dependent. Therefore, secondary mutations strongly depend on the balance between IL-7R and pre-BCR signaling.

Since the proliferative burst is higher following pre-BCR activation than following IL-7R activation, cell survival was analyzed as a function of the presence of mutations downstream of the pre-BCR or the IL-7R, independently of GAL1 expression. Mice with mutations in the IL-7R signaling pathway had a significantly longer median survival than mice with mutations linked to pre-BCR signaling (25 weeks vs. 19 weeks; Figure 7C). All of the mice in the pre-BCR signaling group developed leukemia by 27 weeks, as compared to 60% in the IL-7R signaling group. In addition, the oldest mouse in the IL-7R signaling group to develop leukemia did so at one year of age.

Based on these results, the delay in leukemogenesis in GAL1-deficient mice is related to decreased pre-BCR-dependent proliferative signals and a corresponding increase in IL-7R signaling. Thus, a dysregulation of the balance between pre-BCR and IL-7R signals transmitted by the BM microenvironment can shape the development of pre-leukemic B cells into B-ALL by influencing their cell fate as well as the acquisition of secondary mutations.

## Discussion

The present study aimed to analyze the influence of niche signals, specifically from GAL1, on disease development in two murine models of B-ALL. Our results demonstrated that growth factors produced by the BM microenvironment can not only influence the fate of pre-leukemic cells presenting a first oncogenic hit, but also the acquisition of secondary mutations. GAL1 produced by BM stromal cells favors large pre-B cell proliferation through the activation of pre-BCR signaling ^21,22^. Furthermore, GAL1 can sustain development of both human and mouse pre-BCR^+^ B-ALL ^31^. In murine B cells, the expression of the oncogenic chimeric protein PAX5::ELN identified in human B-ALL induces the development of pre-leukemic cells with features characteristic of cells at the pro-B to large pre-B cell transition, including pre-BCR expression ^11^. Here, the development of pre-leukemic cells was altered in a GAL1-deficient BM microenvironment, not only in terms of relative proportions and proliferative capacity, but also phenotypically. Thus, cells presented an immunophenotype closer to small pre-B cells than cells arrested at the pro-B/large pre-B transition. The scRNAseq analysis of pre-leukemic cells from mice with either a WT background or a GAL1-deficient background confirmed the heterogeneity of these cells, and trajectory inference revealed two distinct pre-leukemic fates, with the pre-leukemic clusters connected to large pre-B cells on one side, and to small pre-B cells on the other. Importantly, in a WT microenvironment, most pre-leukemic cells progressed toward a large pre-B-like cell fate, as confirmed by the sustained pre-BCR expression following B-ALL development. This progression is strongly dependent on GAL1-dependent pre-BCR proliferative signals, as revealed by the pseudo-time analysis. In the absence of GAL1, expression of genes involved in the cell cycle was decreased in favor of differentiation signals (Figure 3L,M). Our previous results showed that IL-7R signaling is reinforced in P5E pre-leukemic cells, whereas pre-BCR differentiation signals are shut down due to the absence of *Blnk* expression ^27^. Our scRNAseq data suggest that, in the absence of GAL1, pre-BCR proliferative signals in large pre-B-like cells are still active and rely on GAL1, in the same way as in normal large pre-B cells ^21^. This conclusion is supported by the parallel decrease in PLCγ2 phosphorylation and in BrdU incorporation (Figure 2F-G).

The absence of GAL1-dependent proliferative signals induced an increase in expression of genes implicated in differentiation, including *Blnk*. In human B-ALL, PAX5::ELN is involved in *Blnk* downregulation regardless of pre-BCR expression ^26^. Evidence presented here demonstrates that GAL1/pre-BCR interactions also control *Blnk* expression, and that this interaction must be arrested to escape the proliferative loop induced in the GAL1^+^ niche. In line with this conclusion, recent results showed that this switch is induced by the lectin function of GAL1, which specifically binds β-galactosides at the termini of glycosylated chains. GAL1 initially strongly binds to LacNac-terminated glycans linked to proteins present at the surface of mesenchymal stromal cells and pre-B cells ^53^. GAL1 binding to the pre-BCR then induces pre-B cell proliferation. However, the interaction with the pre-BCR modifies the carbohydrate-binding affinity of GAL1 to specifically target proteins carrying α2,3-sialylated β-galactosides, triggering a shift between activation and inhibition of proliferative signals ^54^. Furthermore, genes regulated by pre-BCR differentiation signals are also increased on the trajectory toward non-proliferative small pre-B-like cells in the absence of GAL1. This increase confirms that GAL1 impairs pre-BCR differentiation signals, which are consequently likely controlled by ligand-independent tonic signals ^20^. The hypothesis that ligand-independent pre-BCR signals exist in pre-leukemic cells and are implicated in leukemogenesis is supported by the fact that leukemia development was partially impaired in P5EG1^-/-^ compared to P5E animals. In addition, in P5ER2^-/-^ mice, where no pre-BCR is expressed, leukemia development was almost completely blocked (Figure 1G).

GAL1-dependent pre-BCR signaling is not only important for pre-leukemic cell proliferation, but also contributes to pre-leukemic cell quiescence. This result was corroborated by scRNAseq which revealed a decline in cells exhibiting a quiescence signature in the absence of GAL1 (Figure 4B). The biological significance of this observation was validated by the reduced maintenance of pre-leukemic cells from P5E mice following serial transplantation to a GAL1-deficient background. No such reduction was observed upon serial transplantation to a WT background. We previously reported that quiescent pre-leukemic cells in P5E mice are CD117^+ 27^. In line with this characteristic, P5EG1^-/-^ mice presented a decreased proportion of CD117^+^ pre-leukemic cells as compared to P5E mice (Figure 2D). Moreover, upon serial transplantation of P5E pre-leukemic cells in a GAL1-deficient background, their number decreased. Once again, no such decrease in number was seen with serial transplantation on a WT background. In addition, in P5ER2^-/-^ mice, the proportion of pre-leukemic cells was also decreased, strongly limiting leukemia development (Figure 1C,G). Given the absence of pre-BCR in P5ER2^-/-^ pre-leukemic cells, these results clearly suggest that GAL1-dependent pre-BCR signaling contributes to pre-leukemic cell quiescence. Interestingly, in our scRNAseq dataset for normal and pre-leukemic mice, clusters 0 and 4 – where most quiescent cells localized – were both enriched for genes involved in “cell surface interactions at vascular wall” (Figure S2G). These genes not only included *Igll1*, *VpreB1*, *VpreB2* and *VpreB3* coding for the SLC, but also genes involved in adhesion like *Sell*, *Selplg*, *Itgam* or *Sdc4*. This profile further suggests a process dependent on both the pre-BCR and the niche. Tight interactions with the GAL1-expressing niche are particularly important at the large pre-B cell stage ^22,55^ and in pre-BCR^+^ B-ALL ^31^. Furthermore, the deletion of *Ikzf1* (Ikaros) in pre-B cells not only increased their proliferation, but also their adhesion to stromal cells by activating integrins, leading to leukemogenesis ^56^. Accordingly, the expression of *Ikzf1* in large pre-B-like pre-leukemic cells from P5E mice was decreased when GAL1 was expressed (Figure 6M). Altogether, these results confirmed the role of GAL1 in pre-BCR-dependent proliferation and adhesion during leukemogenesis.

Leukemia development in GAL1-deficient animals confirmed the shift observed at the pre-leukemic stage, with a significant increase in BCR^+^ B-ALL blocked at the small pre-B/Immature B cell transition. In WT-background mice, pre-BCR^+^ B-ALL cells were blocked at the pro-B/large pre-B cell transition. Although B-ALL had specific features of tumor cells in contrast to their pre-leukemic counterparts (Figure 6F and S3G), the two populations were relatively similar at the transcriptomic level. In normal small pre-B cells, VpreB and λ5, forming the SLC, are downmodulated ^57^. In contrast, in BCR^+^ B-ALL blocked at the small pre-B cell stage, they were still expressed at high levels, similar to the levels measured in pre-BCR^+^ B-ALL and pre-leukemic cells. Therefore, the block at the pro-B to pre-B cell transition imprinted by the chimeric protein PAX5::ELN is maintained after the acquisition of secondary mutations regardless of pre-BCR or BCR expression.

In the course of leukemia development in P5E mice, secondary mutations of *Ptpn11* and *Jak3* were observed. In the absence of GAL1, pre-BCR activation was decreased in pre-leukemic cells, while STAT5 activation downstream of the IL-7R was increased. In addition, leukemic cells developing in P5EG1^-/-^ mice rarely bore *Ptpn11* mutations, rather they presented a high frequency of *Jak3* mutations, and secondarily of *Jak1, Npm1 and Kmt2d* mutations, all of which play roles downstream of the IL-7R ^47,48^. These results demonstrate that secondary mutational events can be shaped by the microenvironmental context. Chronic stimulation of the pre-BCR by GAL1 combined with signals transmitted by the IL-7R leads to mutations in both pathways. Conversely, in the absence of GAL1-dependent pre-BCR co-signals, mutations are favored downstream of the IL-7R, the other main active pathway at this stage. Although secondary mutations affecting signaling pathways controlled by growth factors most probably relieve leukemic cells of their dependence on their microenvironment, our previous results indicate that BM niches can still contribute to B-ALL maintenance ^31^. Indeed, GAL1 secreted by stromal cells sustains pre-BCR^+^ B-ALL development by activating pre-BCR signals.

Furthermore, despite the presence of *Ptpn11* mutations, the growth of B-ALL originating from P5E mice was impaired in the absence of GAL1 ^31^.

Because of the multiple similarities between mouse and human BM B cell niches, the results presented here are relevant to human disease. Indeed, in addition to the role of GAL1 in pre-BCR signaling in both mouse and human, stromal cells expressing IL7 were identified in human and were found to be very similar to the pro-B cell niche identified in mouse ^16^. Furthermore, cells from B-ALL patients bearing the *PAX5::ELN* translocation are similarly blocked at the pro-B to pre-B cell transition and present mutations of the ERK and IL-7R signaling pathways ^11,26^.

In conclusion, our findings demonstrate that leukemogenesis can be influenced by growth factors produced by BM niches, and that variations in the BM milieu after the acquisition of the first oncogenic hit can guide the (pre)-leukemic cell fate and the ensuing secondary events. Therefore, the accumulation of dysregulated cells that are unable to escape their niche, and are consequently exposed to chronic signals, represents a pivotal step in leukemogenesis. This phenomenon has been observed in the *PAX5::ELN* model and can be extrapolated to other driver mutations. The *BCR::ABL1* translocation has been identified in several B-ALL subtypes exhibiting differentiation arrests from the early pro-B to the pre-B cell stage with stage-dependent secondary mutations ^31,58,59^. How the secondary mutations are selected in this particular case remains to be elucidated. The associated chronic signals are also important for the resistance of leukemic cells to treatment, since blocking pre-BCR-dependent signaling while pathways induced by driver mutations are also inhibited decreases relapse rates ^31^. Therefore, identifying growth factors that have the ability to fuel leukemogenesis and sustain leukemia development could lead to the emergence of adjuvant strategies that deplete treatment-resistant leukemic cells by blocking the supply of growth factors. These strategies could be used alongside chemotherapy.

## Methods

### Mice

PAX5::ELN (P5E) mice generated by AA Khamlishi ^11^ were crossed with *Rag2*^-/-^ ^28^ and *Lgals1*-/-mice ^60^ to obtain *PAX5::ELN/Rag2*^-/-^ (P5ER2^-/-^) and *PAX5::ELN/Lgals1*^-/-^ (P5EG1^-/-^) mice, respectively. Mice were backcrossed for at least 8 generations onto the C57Bl/6J background (obtained from Janvier Laboratories, France). Both males and females were used and housed under specific pathogen-free conditions. Pre-leukemic mice were analyzed at 5-7 week-old, age at which secondary mutations are not detected ^11^. For the analysis of leukemic development, mice were monitored until mice reached the limit point. Mice were handled in accordance with the French Guidelines for animal handling and the EU Directive 2010/63/EU.

### Serial transplantation assays

P5E^Tg+/-^ mice under the C57Bl/6J background (CD45.2 haplotype) were intercrossed with B6-Ly5.1 mice (CD45.1 haplotype; Charles River Laboratories, France) to obtain CD45.2^+^CD45.1^+^ hematopoietic cells. Donor pre-leukemic cells (B220^lo^CD19^+^) were sorted from these mice on a FACSAria III (BD Biosciences) and 8x10^5^ cells were transplanted i.v. to both 6–9-week-old C57Bl/6J and *Lgals1*^-/-^ recipients. For the secondary and tertiary transplants, 5x10^6^ total BM cells from the primary transplants into C57Bl/6J and *Lgals1*-/- were engrafted i.v. to 6–9-week-old C57Bl/6J and *Lgals1*^-/-^ recipients, respectively.

### Flow cytometry

Femurs and tibias were flushed with FACS Buffer (PBS 1X, 0.1% FCS, 2 mM EDTA). Red blood cells were lysed (Gibco ACK Lysing Buffer) and mouse B cells stained with antibodies for 20 min at 4 °C. Dead cells were excluded using Sytox Blue, LIVE/DEAD Fixable Aqua Dead Cell Stain (Thermo Fisher) or LIVE/DEAD Fixable Far Red Cell Stain (Thermo Fisher). Intracellular staining was performed using a BD Cytofix/Cytoperm Fixation/Permeablization kit (BD Biosciences). Pre-leukemic cells were defined as being CD19^+^B220^lo^CD45^lo^. B cell subsets were defined as described previously ^14^: pro-B: CD19^+^B220^+^Igκ/λ^-^CD23^-^BP1^-^CD2^-^CD117^+^; large pre-B: CD19^+^B220^+^Igκ/λ^-^CD23^-^BP1^+^CD2^+^CD117^-^ and large FSC; small pre-B: CD19^+^B220^+^Igκ/λ^-^CD23^-^BP1^+/-^CD2^+^CD117^-^ and small FSC; Immature B cells: CD19^+^B220^+^Igκ/λ^+^CD23^-^; Recirculating B cells: CD19^+^B220^+^Igκ/λ^+^CD23^+^.

FACS acquisitions were performed on a LSRII and a FortessaX20 (BD Boisciences). Data were analyzed using DiVa v9.0.1 (BD Biosciences) or FlowJo Version 10 (TreeStar) software.

For the analysis of phospho-proteins, femurs and tibias were flushed with cold RPMI 2%FCS containing protease/phosphatase inhibitors. Surface staining was performed on ice. Intracellular staining was performed using anti pPLCγ2 (clone K86-689.37) and anti-pSTAT5 (clone 47/Stat5) antibodies with the PerFix EXPOSE Kit (Beckman Coulter), following the manufacturer instructions.

### *In vivo* BrdU incorporation

Mice at 5-7 weeks are injected i.p. with 1 mg/mL BrdU (BrdU Flow Kit, BD Biosciences). Femurs and tibias were dissected 16 hours after injection and cells were stained following the manufacturer instructions. Briefly, after surface staining, cells were fixed and permeabilized with Cytofix/Cytoperm buffer and washed twice with Perm/Wash buffer before treatment with 300 µg/mL DNase I 1 h at 37 °C. After washing the cells in Perm/Wash buffer, cells were resuspended in Perm/Wash buffer containing the anti-BrdU FITC antibody and incubated 20 min at RT. The samples were acquired by flow cytometry.

### Single cell RNAseq analysis

CD19^+^ B cells were sorted from 5-week-old WT, G1^-/-^, P5E and P5EG1^-/-^ mice. Leukemic samples obtained from the BM of P5E and P5EG1^-/-^ were stored in nitrogen and thawed to be sorted based on the expression of CD19. Samples were tagged individually with anti-CD45 conjugated to custom barcodes for the first dataset and with the anti-H2/CD45 TotalSeq-C (Biolegend) for the second dataset.

The library of the first dataset was generated with the Chromium Next GEM Single Cell 5’ Library and Gel Bead Kit v1.1 with Feature barcode for cell surface protein & repertoire (10X Genomics), and the library of the second dataset was generated with the Chromium Next GEM Single Cell 5’ Reagent Kits v2 with Feature barcode for cell surface protein & repertoire. This protocol was adapted for single index to be in similar conditions as the first dataset.

Fastq raw files were processed using the Multi pipeline from Cell Ranger v7.1.0 (reads alignment to the mm10 murine genome, barcode sample assignment, VDJ profiling, filtering, barcode counting and unique molecular identifier (UMI) counting). scRNAseq pre-processing was performed using Seurat v4.1.3. The demultiplexing of hash tag oligonucleotides was performed using the HTOdemux function with a manual curation. Cells were filtered based on the following parameters for both datasets: feature counts between 750 and 5500; reads related to mitochondrial genes below 10%; ribosomal reads above 10%. For the leukemic samples, B-ALL cells expressing the dominant IgH clonotype were kept for further analysis. Number of cells after filtering in dataset1: WT-1: 1068 cells; WT-2: 1177 cells; G1ko-1: 1217 cells; G1ko-2: 1252 cells; preleuk P5E-1: 1126 cells; preleuk P5E-2: 1173 cells; preleuk P5EG1-1: 1160 cells; preleuk P5EG1-2: 1229 cells; preleuk P5EG1-3: 1270 cells; preleuk P5EG1-4: 1272 cells. Number of cells after filtering in dataset2: Leuk P5E-1: 488 cells; Leuk P5E-2: 1005 cells; Leuk P5E-3: 1056 cells; Leuk P5EG1-1: 1153 cells; Leuk P5EG1-2: 1277 cells; Leuk P5EG1-3: 1199 cells; Preleuk P5E: 1382 cells.

Data were normalized using the NormalizeData function and the UMAP clustering was performed as follow: the 2000 most variable genes were identified with the FindVariableFeature function. As immunoglobulin genes are the most variable genes, these genes were removed before computing the PCA to avoid a clustering mainly influenced by their expression. The scaling was performed with the ScaleData function and the PCA with the RunPCA function. The UMAP and the clustering were generated with the RunUMAP, FindNeighbors and FindClusters functions with the 10 first PCA dimensions and the neighbor parameter set to 45.

Integration of pre-leukemic cells from the first dataset with pre-leukemic and leukemic cells from the second dataset was performed with the FindIntegrationAnchors and IntegrateData functions with the reduction and number of neighbors set to “rpca” and 45 respectively. Integrated expression data was processed as previously described (ScaleData, RunPCA, RunUMAP, FindNeighbors, FindClusters functions) with 20 PCA dimensions used for UMAP and neighbors computing.

Differentially expressed genes were identified with the FindMarkers function with the default parameters. The repertoire analysis was performed using the scRepertoire package v1.11.0 ^61^.

Cell cycle analysis was performed using the CellCycleScoring function. S and G2M signatures from Seurat package were previously converted to mouse names with the biomaRt package (v2.54.1). Signature scores were generated by using the AddModuleScore function. Scores used to initially identify the differentiating B cell subsets, the IL-7R signaling, and the pre-BCR proliferation or differentiation pathways on the first dataset were generated based on previous studies and phenotypic knowledge ^14,19,32,62,63^. After attribution of the B cell subset to the various clusters, more specific signatures were generated from the top 50 inter-cluster DEGs using only the WT cells of the dataset.

Scores for the quiescent and proliferating states were generated using differentially expressed genes from bulk RNAseq data of proliferating and quiescent cells from P5E pre-leukemic cells ^27^. Cells were tagged as “intermediate” if both scores were < 0. All other cells were considered as “proliferating” or “quiescent” according to the signature giving the maximum score in each cell.

Enrichment analysis was performed using Enrichr ^64^. Enrichments shown are from the Reactome and MSigDB Hallmark databases. For each cluster, the genes differentially expressed with an adjusted p-value<0.05 and a fold change > 1.3 were used.

Trajectory analysis was performed with the monocle3 package v1.4.23 using the learn_graph function. Cells were order with the order_cells function by choosing the extremity of the pro-B cluster as starting point. Resulting pseudotime values were rescaled to synchronize the two converging branches.

Estimation of gene expression profiles across pseudotime was computed with the fitGAM function from the tradeSeq package v1.4.

### Secondary mutation analysis

Targeted sequencing was performed on *Jak3* (exons 13 and 15), *Kras* (exons 2 and 3), *Nras* (exons 2 and 3) and *Ptpn11* (exons 3 and 11) on genomic DNA from the bone marrow of 18 P5E and 12 P5EG1^-/-^ mice as described previously ^11^. The library was sequenced using a MiSeq sequencer (Illumina, San Diego, USA) and MiSeq Reagent kit V2 (paired-end sequencing 2 x 150 cycles). Alignment and variant calling were performed using NextGene software (SoftGenetics, State College, USA).

Whole exome sequencing was performed using genomic DNA extracted from the BM of leukemic P5E and P5EG1^-/-^ mice. The samples were prepared by Macrogen according to an Agilent SureSelect Target Enrichment Kit preparation guide. The region of interest was enriched using a SureSelect Target Enrichment workflow. The libraries were sequenced with Illumina HiSeq 2000/2500/4000 sequencer. Sequences were mapped to a reference genome from a pre-leukemic P5E mouse using BWA software program. Variant discovery was performed with Genome Analysis toolkit or GATK software. Variants were annotated and effects predicted using the SnpEff tool.

### Quantification and statistical analysis

Statistical analysis was performed using GraphPad Prism v10 software. Data are presented as mean values with error bars representing the SEM. Sample distribution was assessed using a Shapiro-Wilk normality test. Samples were compared using a two-tailed unpaired t-test or a Mann-Whitney test when normality was not reached.

The frequencies of pre-BCR^+^ vs. BCR^+^ B-ALL from P5E and P5EG1^-/-^ mice were compared using a Fisher’s exact test from a contingency table. *, p < 0.05; **, p < 0.01; ***, p < 0.001; ****, p<0.0001.

### Data availability

The accession numbers for the scRNAseq datasets reported in the study are available in the GEO database under accession number GSEXXXXX and GSEXXXXX

**Table 1.**

| REAGENT OR RESSOURCE | SOURCE | IDENTIFER |
| --- | --- | --- |
| **Antibodies** |  |  |
| B220 Alexa700 clone RA3-6B2 | eBioscience | Cat# 56-0452-82 |
| B220 APC clone RA3-6B2 | eBioscience | Cat# 17-0452-83 |
| BP1 BV786 clone BP-1 | BD Bioscience | Cat# 740882 |
| CD71 BV605 clone C2 | BD Bioscience | Cat# 563013 |
| CD117 BV605 clone 2B8 | BD Bioscience | Cat# 563146 |
| CD117 PE clone 2B8 | eBioscience | Cat# 12-1171-83 |
| CD127 APC clone A7RB4 | Bio Legend | Cat# 135020 |
| CD19 BV605 clone 1D3 | BD Bioscience | Cat# 563148 |
| CD19 PE-Cy5.5 clone 1D3 | eBioscience | Cat# 15-0193-83 |
| CD2 FITC clone RM2-5 | eBioscience | Cat# 11-0021-82 |
| CD2 PE-Cy7 clone RM2-5 | Bio Legend | Cat# 100114 |
| CD23 BV711 clone B3B4 | BD Bioscience | Cat# 563987 |
| CD25 biotin clone PC61 | BD Bioscience | Cat# 13-0251-85 |
| CD45 PE-Cy5.5 clone 30-F11 | eBioscience | Cat# 35-0451-82 |
| CD45.1 PE-CF594 clone A20 | BD Bioscience | Cat# 562452 |
| CD45.1 PE-Cy5 clone A20 | eBioscience | Cat# 15-0453-82 |
| CD45.2 biotin clone 104 | eBioscience | Cat# 13-0454-85 |
| CD45.2 FITC clone 104 | eBioscience | Cat# 11-0454-85 |
| CD93 PE-Cy7 clone AA4.1 | eBioscience | Cat# 25-5892-82 |
| Gr1 Alexa700 clone RB6-8B5 | BD Bioscience | Cat# 557979 |
| Gr1 PE-Cy5 clone RB6-8B5 | eBioscience | Cat# 15-5931-83 |
| Igκ BV421 clone 187.1 | BD Bioscience | Cat# 562888 |
| Igλ BV421 clone R26-46 | BD Bioscience | Cat# 744523 |
| Igκ FITC clone 187.1 | BD Bioscience | Cat# 550003 |
| Igλ FITC clone R26-46 | BD Bioscience | Cat# 553434 |
| IgM APC polyclonal | Southern Biotechnologies | Cat# 1021-31 |
| IgM BV421 clone R6-60.2 | BD Bioscience | Cat# 562595 |
| IgM FITC polyclonal | Southern Biotechnologies | Cat# 1021-02 |
| IgM PE polyclonal | Southern Biotechnologies | Cat# 1021-09 |
| IgM PE-CF594 clone R6-60.2 | BD Bioscience | Cat# 562565 |
| Ki67 V450 | BD Bioscience | Cat# 561281 |
| Pre-BCR biotin clone SLC156 | BD Bioscience | Cat# 551863 |
| Streptavidin PE-Cy5 | BD Bioscience | Cat# 554062 |
| Streptavidin PE-Cy5.5 | eBioscience | Cat# E032716 |
| Streptavidin APC-Cy7 | BD Bioscience | Cat# 554063 |
| Streptavidin BV421 | BD Bioscience | Cat# 563259 |
| Anti-Stat5 (pY694) Alexa Fluor 647 clone 47/Stat5(pY694) | BD Bioscience | Cat# 612599 |
| Anti-PLCy2 APC (pY759) Alexa fluor 647, clone K86-689.37 | BD Bioscience | Cat# 558498 |
| **Chemicals, Peptides, and Recombinant Proteins** |  |  |
| SYTOX Blue Nucleic Acid Stain | ThermoFisher | Cat# S11348 |
| LIVE/DEAD Fixable Aqua Dead Cell Stain Kit | ThermoFisher | Cat# L34957 |
| 7-AAD | ThermoFisher | Cat#A1310 |
| TOPRO 3 | Invitrogen | Cat# R37113 |
| BrdU Flow Kits | BD Pharmingen | Cat#559619 |
| PEI cocktail |  | **N/A** |
| **Biological samples** |  |  |
| Primary murine leukemic and pre-leukemic cells from P5E mice | This paper | N/A |
| Primary murine leukemic and pre-leukemic cells from P5EG1KO mice | This paper | N/A |
| **Critical Commercial Assays** |  |  |
| 1X RBC Lysis Buffer | eBioscience | Cat# 00-4333-57 |
| ACK Lysing Buffer | Gibco | Cat# A1049201 |
| PerFix EXPOSE (Phospho-Epitopes Exposure kit) | Beckman and coulter | Cat# B26976 |
| **Experimental Models: Organisms/Strains** |  |  |
| Mouse: C57BL/6J (CD45.2) | Charles River | N/A |
| Mouse: Ly5.1 (CD45.1) | Charles River | B6.SJL-PtprcaPepcb/BoyCrl |
| Mouse: Pax5-ELN Tg<sup>+/-</sup> | Jamrog et al. 2018 | N/A |
| Mouse: Lgals1<sup>-/-</sup> | Poirier and Robertson, 1993 | N/A |
| Mouse: Pax5-ELN Tg<sup>+/-</sup> Lgals1<sup>-/-</sup> | This paper | N/A |
| **Software and algorithms** |  |  |
| DIVA version 9.0.1 | BD Biosciences | http://www.bdbiosciences.com/ |
| FlowJo version 10 | TreeStar | https://www.flowjo.com/ |
| Prism version 5.3 | GraphPad | https://www.graphpad.com/ |

## Acknowledgements

We are grateful to the core flow cytometry and the animal facility of the CRCM for providing supportive help. We are also grateful to the ARCHE and flow cytometry facilities of the UAR Biosit in Rennes. We thank the members of the CB2M (Computational Biology, Biostatistics and Modeling) group at CIML for their expert help with data analysis and in particular Lionel Spinelli for our fruitful discussions.

This work was supported by the Institut National du Cancer (INCa-2020-096), the Ligue Nationale Contre le Cancer (#ELN2020), the ADHO association and Rennes Métropole. M.C.D. was the recipient of PhD grants from la Ligue Nationale contre le Cancer (IP/SC-16060) and the ARC Foundation; J.P. from the Région PACA/Inserm/Innate Pharma and from La Ligue Nationale contre le Cancer (#TDCU19117); M.O from the Région Bretagne/Ligue Contre le Cancer.

