## Supplementary material for "Signals from the bone marrow B cell niches shape pre-leukemic fate in murine B cell acute lymphoblastic leukemia"

* co-second authors

§ senior co-authors

**SUPPLEMENTARY INFORMATION**

**Supplementary figure legends**

**Figure S1.** Related to Figure 2.

(**A**) Percentage of CD2$^+$Igκ/λ$^-$ pre-B cells among CD19$^+$B220$^+$ B cells for WT (n=14), G1$^{-/-}$ (n=13), P5E (n=15), and P5EG1$^{-/-}$ (n=20) mice. (**B**) Percentage of Immature B cells, B220$^+$CD19$^+$Igκ/λ$^+$CD23$^-$, and (**C**) of recirculating B cells, B220$^+$CD19$^+$Igκ/λ$^+$CD23$^+$, in WT (n=20), G1$^{-/-}$ (n=19), P5E (n=22), and P5EG1$^{-/-}$ (n=30) mice. For parametric data, statistical significance was tested using a t-test. *: p-val<0.05; **: p-val<0.01; ***: p-val<0.001; ****: p-val<0.0001. For non-parametric data, statistical significance was tested using a Mann-Whitney test. #: p-val<0.05; ##: p-val<0.01; ###: p-val<0.001; ####: p-val<0.0001.

**Figure S2.** Related to Figure 3.

(**A**) and (**B**) Scores generated for signatures of differentiating B cells or for signatures of the IL-7R pathway and pre-BCR proliferating / differentiating pathways. Scores are represented as a heatmap on the UMAP space for WT and G1$^{-/-}$ mice, and in a violin plot for each cluster. (**C**) Each cell on the UMAP was tagged with its cell cycle signature by defining a score based on genes characteristic of the S and G2M phases. The overall distribution of B cells for the four genotypes is represented in the UMAP space. (**D**) Expression of *Iglc1*, *Iglc2* and *Iglc3* in the UMAP space. (**E**) Intra-cluster DEG for WT vs. GAL1$^{-/-}$ mice for the large pre-B cell clusters (clusters 7-8-9) were defined, and the enrichment of the genes significantly upregulated by WT cells was analyzed. (**F**) The IgH repertoire was determined for each cell in the dataset. Ig repertoire clonality was based on IgH expression. (**G**) and (**H**) Reactome and MSigDB enrichment analysis of inter-cluster DEG for the clusters indicated in the panels. (**I**) Proportion of quiescent cells in subclusters s0 to s5 and in cluster 6 for P5E and P5EG1$^{-/-}$ cells.

**Figure S3.** Related to Figure 6

(**A**) The IgH and IgL repertoires were determined for each cell in the dataset. Ig repertoire clonality was based on IgH expression. Cells were split by their decreasing occupancy in the repertoire space. (**B**)

Proportion of cells identified according to their repertoire profile for each sample. (**C**) Proportion of the different IgL clonotypes identified in each B-ALL sample represented by a monoclonal IgH clonotype. The most represented CDR3 sequences and the number of clonotypes and of $IgL^+$ cells are shown. (**D**) Expression of genes coding for λ5 (*Igll1*) and VpreB (*Vpreb1, Vpreb2, Vpreb3*) in the leukemic and pre-leukemic samples. (**E**) Distribution in the UMAP space of cells for each of the leukemic and pre-leukemic samples. (**F**) Heatmap showing the expression of inter-cluster DEG obtained from the pre-leukemic clusters (left panel) and subclusters (right panel) of the dataset shown in Figure 3, for cells present in the clusters of the leukemic/pre-leukemic integrated UMAP from Figure 6. (**G**) Expression of *S100a4* and *s100a6* by leukemic and pre-leukemic cells.

**Figure S1**

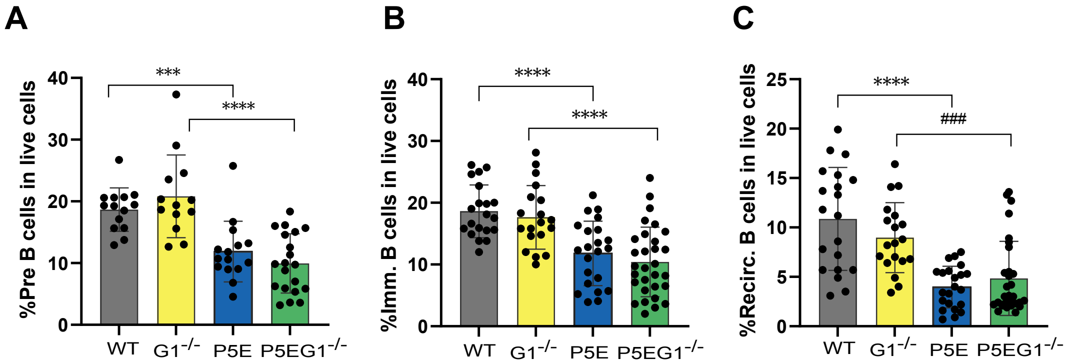
A
%Pre B cells in live cells
0
10
20
30
40
***
****
WT
G1-/-
P5E
P5EG1-/-
B
%Imm. B cells in live cells
0
10
20
30
40
****
****
WT
G1-/-
P5E
P5EG1-/-
C
%Recirc. B cells in live cells
0
5
10
15
20
25
****
###
WT
G1-/-
P5E
P5EG1-/-

### Figure S2

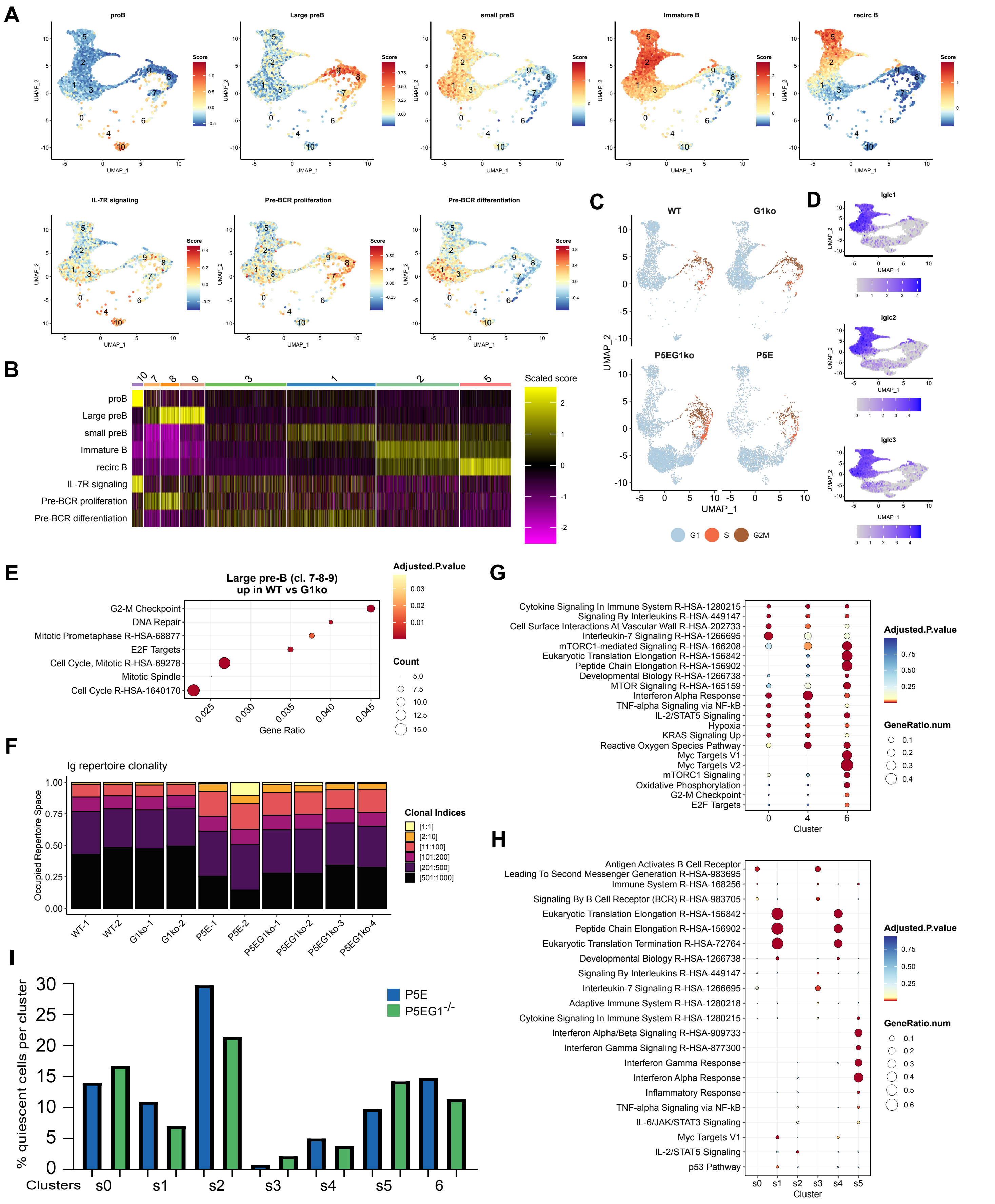

A
proB
Large preB
small preB
Immature B
recirc B
IL-7R signaling
Pre-BCR proliferation
Pre-BCR differentiation
Score
UMAP_1
UMAP_2
B
Scaled score
proB
Large preB
small preB
Immature B
recirc B
IL-7R signaling
Pre-BCR proliferation
Pre-BCR differentiation
C
WT
G1ko
P5EG1ko
P5E
UMAP_1
UMAP_2
G1
S
G2M
D
Iglc1
Iglc2
Iglc3
E
Large pre-B (cl. 7-8-9)
up in WT vs G1ko
G2-M Checkpoint
DNA Repair
Mitotic Prometaphase R-HSA-68877
E2F Targets
Cell Cycle, Mitotic R-HSA-69278
Mitotic Spindle
Cell Cycle R-HSA-1640170
Gene Ratio
Adjusted.P.value
Count
F
Ig repertoire clonality
Occupied Repertoire Space
WT-1
WT-2
G1ko-1
G1ko-2
P5E-1
P5E-2
P5EG1ko-1
P5EG1ko-2
P5EG1ko-3
P5EG1ko-4
Clonal Indices
[1:1]
[2:10]
[11:100]
[101:200]
[201:500]
[501:1000]
G
Cytokine Signaling In Immune System R-HSA-1280215
Signaling By Interleukins R-HSA-449147
Cell Surface Interactions At Vascular Wall R-HSA-202733
Interleukin-7 Signaling R-HSA-1266695
mTORC1-mediated Signaling R-HSA-166208
Eukaryotic Translation Elongation R-HSA-156842
Peptide Chain Elongation R-HSA-156902
Developmental Biology R-HSA-1266738
MTOR Signaling R-HSA-165159
Interferon Alpha Response
TNF-alpha Signaling via NF-kB
IL-2/STAT5 Signaling
Hypoxia
KRAS Signaling Up
Reactive Oxygen Species Pathway
Myc Targets V1
Myc Targets V2
mTORC1 Signaling
Oxidative Phosphorylation
G2-M Checkpoint
E2F Targets
Cluster
Adjusted.P.value
GeneRatio.num
H
Antigen Activates B Cell Receptor
Leading To Second Messenger Generation R-HSA-983695
Immune System R-HSA-168256
Signaling By B Cell Receptor (BCR) R-HSA-983705
Eukaryotic Translation Elongation R-HSA-156842
Peptide Chain Elongation R-HSA-156902
Eukaryotic Translation Termination R-HSA-72764
Developmental Biology R-HSA-1266738
Signaling By Interleukins R-HSA-449147
Interleukin-7 Signaling R-HSA-1266695
Adaptive Immune System R-HSA-1280218
Cytokine Signaling In Immune System R-HSA-1280215
Interferon Alpha/Beta Signaling R-HSA-909733
Interferon Gamma Signaling R-HSA-877300
Interferon Gamma Response
Interferon Alpha Response
Inflammatory Response
TNF-alpha Signaling via NF-kB
IL-6/JAK/STAT3 Signaling
Myc Targets V1
IL-2/STAT5 Signaling
p53 Pathway
Cluster
Adjusted.P.value
GeneRatio.num
I
% quiescent cells per cluster
P5E
P5EG1-/-
Clusters
s0
s1
s2
s3
s4
s5
6

### Figure S3

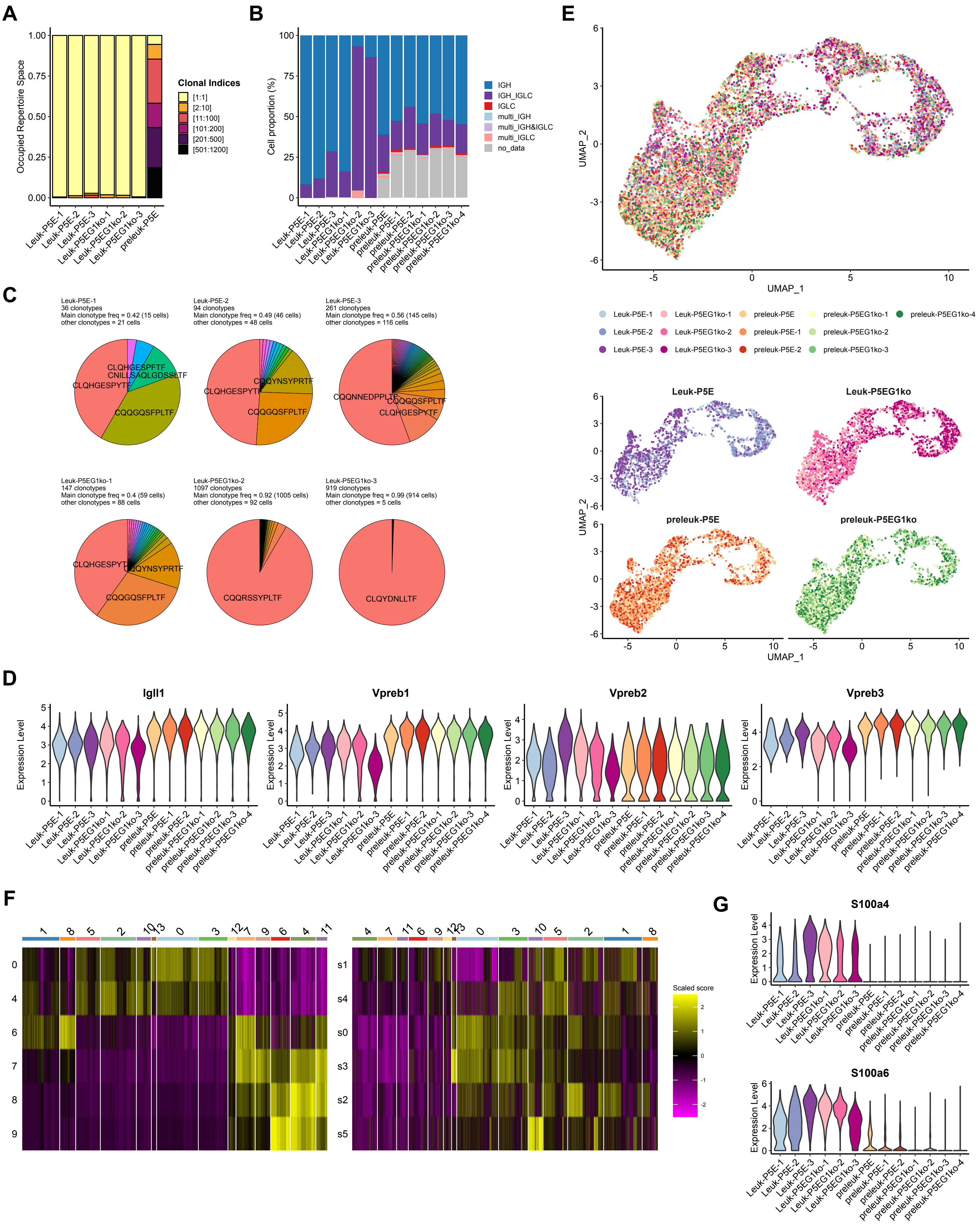
A
Occupied Repertoire Space
Clonal Indices
[1:1]
[2:10]
[11:100]
[101:200]
[201:500]
[501:1200]
Leuk-P5E-1
Leuk-P5E-2
Leuk-P5E-3
Leuk-P5EG1ko-1
Leuk-P5EG1ko-2
Leuk-P5EG1ko-3
preleuk-P5E
B
Cell proportion (%)
IGH
IGH_IGLC
IGLC
multi_IGH
multi_IGH&IGLC
multi_IGLC
no_data
preleuk-P5E-1
preleuk-P5E-2
preleuk-P5EG1ko-1
preleuk-P5EG1ko-2
preleuk-P5EG1ko-3
preleuk-P5EG1ko-4
C
Leuk-P5E-1
36 clonotypes
Main clonotype freq = 0.42 (15 cells)
other clonotypes = 21 cells
Leuk-P5E-2
94 clonotypes
Main clonotype freq = 0.49 (46 cells)
other clonotypes = 48 cells
Leuk-P5E-3
261 clonotypes
Main clonotype freq = 0.56 (145 cells)
other clonotypes = 116 cells
CLQHGESPFTF
CNILLSAQLGDSSLTF
CLQHGESPYTF
CQQGQSFPLTF
CQQYNSYPRTF
CQQNNEDPPLTF
Leuk-P5EG1ko-1
147 clonotypes
Main clonotype freq = 0.4 (59 cells)
other clonotypes = 88 cells
Leuk-P5EG1ko-2
1097 clonotypes
Main clonotype freq = 0.92 (1005 cells)
other clonotypes = 92 cells
Leuk-P5EG1ko-3
919 clonotypes
Main clonotype freq = 0.99 (914 cells)
other clonotypes = 5 cells
CQQRSSYPLTF
CLQYDNLLTF
D
Igll1
Vpreb1
Vpreb2
Vpreb3
Expression Level
E
UMAP_1
UMAP_2
Leuk-P5E
Leuk-P5EG1ko
preleuk-P5E
preleuk-P5EG1ko
F
Scaled score
s1
s4
s0
s3
s2
s5
G
S100a4
S100a6
